# Epi-MEIF, a flexible and efficient method for detection of high order epistatic interactions from complex phenotypic traits

**DOI:** 10.1101/2021.12.21.473474

**Authors:** Saswati Saha, Laurent Perrin, Laurence Röder, Christine Brun, Lionel Spinelli

## Abstract

Understanding the relationship between genetic variations and variations in complex and quantitative phenotypes remains an ongoing challenge. While Genome-wide association studies (GWAS) have become a vital tool for identifying single-locus associations, we lack methods for identifying epistatic interactions. In this article, we propose a novel method for high-order epistasis detection using mixed effect conditional inference forest (*epiMEIF*). The *epiMEIF* model is fitted on a group of potential causal SNPs and the tree structure in the forest facilitates the identification of n-way interactions between the SNPs. Additional testing strategies further improve the robustness of the method. We demonstrate its ability to detect true n-way interactions via extensive simulations in both cross-sectional and longitudinal synthetic datasets. This is further illustrated in an application to reveal epistatic interactions from natural variations of cardiac traits in flies (Drosophila). Overall, the method provides a generalized way to identify high order interactions from any GWAS data, thereby greatly improving the detection of the genetic architecture of complex phenotypes.

## 1. Introduction

Over the past few decades, there has been a growing interest in phenotype to genotype association and Genome-Wide Association Studies (GWAS) have proven to be ‘the standard tool’ for identifying these associations^1,2^. GWAS have attained tremendous success in identifying causal variants that exhibit independent, additive, and cumulative effects on the investigated phenotype trait^3^. However, testing for associations *via* a single-locus test is an oversimplified approach to tackle the complexity of underlying biological mechanisms^4^. Complex phenotypes and their variations within a population are speculated to be caused by multiple genetic variations and their interactive effects, which are referred to as *epistatic interactions*^5–8^. Nonetheless, the exhaustive evaluation of all possible *epistatic interactions* among millions of single nucleotide polymorphisms (SNPs) raises several issues, otherwise known as the ‘curse of dimensionality’. Indeed, due to the exponential complexity involved in the higher-order exhaustive search algorithms, they are not applicable to large datasets. To address the above issues, several parametric modelling approaches^9,10^, machine learning algorithms^8,11^, and combinatorial optimizations^12,13^ have been explored. But they are exclusively designed/used for detecting binary or higher-order interactions in case-control studies. There exist some approaches^14–17^ that can detect pairwise interactions in crosssectional studies with quantitative traits but they are simply not scalable to higher-order interactions. Moreover, current methods for detecting epistasis do not address the complexities and challenges involved in longitudinal datasets that allow studying the natural trajectory of traits and/or disease progression.

In this regard, we propose a novel approach of epistasis detection, called *epiMEIF*, using mixed-effect conditional inference forest (*MEIF*). **The goal of our approach is to reveal higher-order epistatic interactions of genetic variants associated with the variation of complex quantitative phenotypes, primarily from cross-sectional and longitudinal GWAS datasets.** The mixed effect conditional inference forest (*MEIF*) algorithm we propose is a combination of mixed effects model and conditional inference forest (see Figure 1 and Methods, for more details). Recursive partitioning approaches or tree-based algorithms like random forest have already proven to be effective for detecting the genetic loci and their interactions that impact the phenotypic outcome in case-control studies^7,8,18,19^. Nevertheless, *MEIF* is particularly beneficial over existing methods because it i) **demonstrates how tree-based algorithms can be adapted for detecting higher-order interactions from cross-sectional and longitudinal datasets using conditional inference forest (cforest), ii) can handle ‘large’ datasets (as much as 10000 markers)** and iii) simultaneously **account for missing/censored genome data using cforest,** and **other confounding factors like population structure in the GWAS datasets using mixed effects model.** Subsequent to identifying interactions from MEIF, we propose to validate the epistatic interactions using two statistical testing approaches: Max T-test and anova test (see Methods, for more details). Overall, *epiMEIF* (epistasis detection from MEIF along with the additional testing strategies), provides a generalized way to obtain genetic variants and their higher order interactions from any GWAS data.

**Figure 1:**
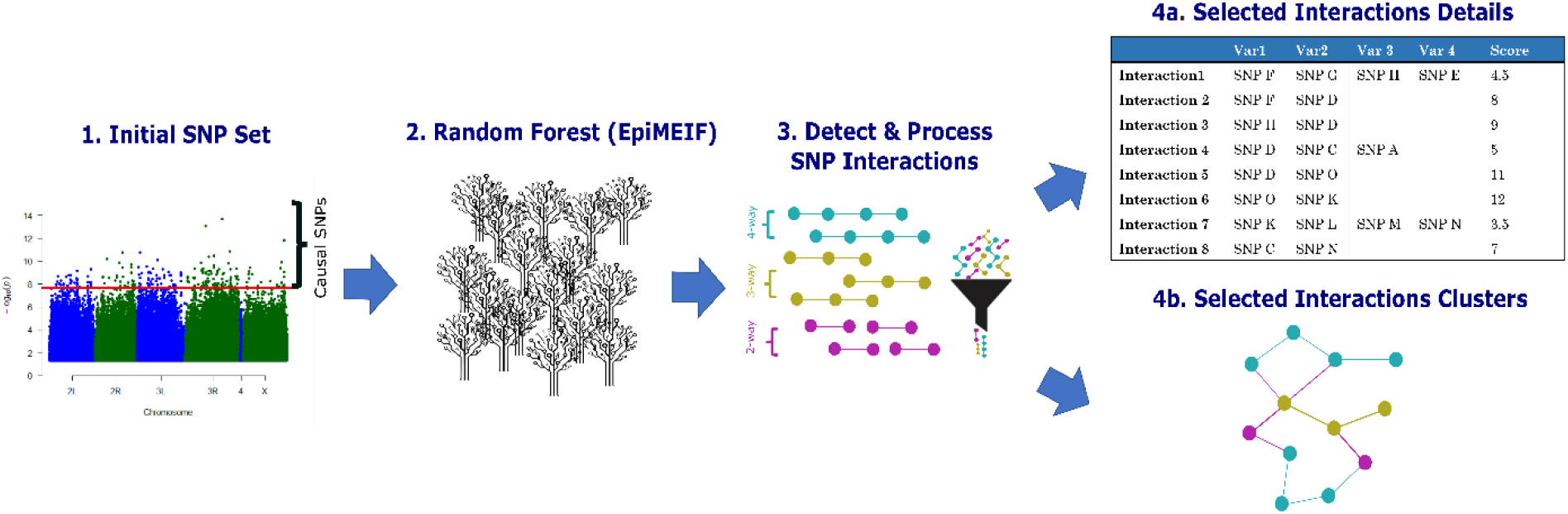
Global overview of *epiMEIF*. 1) The method begins with a set of causal SNPs obtained from a single-locus association test (*via* LMM). SNP markers that have a nominally significant effect on the phenotype are selected. 2) The mixed effect conditional inference forest (MEIF) is applied on the phenotype, selected SNPs and other additional covariates that might explain the variability of the phenotype. 3) Identified SNP interactions from MEIF are subjected to additional statistical testing (anova and Max-T test) that helps in filtering the stable interactions. 4) The final set of interactions can be visualized in two ways: a) the interaction score table that captures the different sets of interaction and their associated score from epiMEIF and b) as an interaction network where the nodes denote the different variants, and the edges denote the interactions from *epiMEIF*. Different order interactions are illustrated by different coloured edges in the network.

To the best of our knowledge, alternative approach that can detect higher-order interactions in both cross-sectional and longitudinal datasets does not exist, therefore preventing us from evaluating our method *via* benchmarking. Hence, to evaluate our method, we applied it to an extensive simulation study and illustrated the power performance of the method under several practically relevant scenarios. We then analysed its performances on a real dataset: natural variation of heart period in young flies (Drosophila) and heart period during aging in Drosophila. We evaluated the method’s performance based on its ability to replicate in an independent cohort and to recover previously published genetic data associated with cardiac functions. We demonstrate that the obtained networks of statistical interactions provide insightful information with respect to cardiomyocytes structure and functions. These analyses illustrate the high performance of the method to identify higher-order interactions from both cross-sectional and longitudinal data. The proposed statistical methods are implemented in R and the source codes can be found in the Github repository (https://github.com/TAGC-NetworkBiology/epiMEIF).

## 2. Results

### 2.1 Overview of computational methods

In this article, we propose a new method for epistasis detection in large scale association studies with complex genetic traits, using mixed effect conditional inference forest (*epiMEIF*). The random forest technique is a tree-based predictive method, which produces a series of classification/regression trees using a large set of predictor variables. It has so far been used to discover the SNPs and their interactions that are most predictive of the disease status in large-scale case-control association studies^7,8,20^. Conditional inference forests (cforest) are an alternative to random forests that are particularly useful for complex case studies with missing or censored data. Using cforest, we have extended the random forest framework so that it is applicable for any type of complex phenotypes — cross-sectional or longitudinal — with missing genotype data, and exploited the tree structure to detect higher-order SNP interactions.

The method begins with fitting the MEIF model on a set of causal SNPs (and additional covariates, if any) and exploits the tree structure in cforest to compute the SNP Interaction matrix (*I*). It measures the strength of each interaction set based on their frequency of occurrence in the same path of the forest. Thereafter, the interactions in *I* are validated using additional testing strategies (Max-T test or anova test) leading to the final SNP Interaction matrix (*F*). These independent testing strategies increase the reliability of the final interactions and help to get rid of false positives, if any. Finally, the SNP Interaction matrix (*F*) helps to build the statistical epistatic network of genetic variants (see Methods). The complete workflow is illustrated in Figure 1. Seldom, the tree-based algorithms are criticised to be biased towards variants with high marginal effect. To address that caveat, we developed a weighted adaptation of mixed effect conditional inference forest (Weighted *epiMEIF*), where the probability of selecting the variant during the root node construction of the tree depends on the significance of the predictor variable from the single-locus association; higher the significance, lower the sampling weight (see Methods, for more details).

### 2.2 Application on Cross-sectional Data

#### 2.2.1 Analysis on Simulated Data

We evaluated the statistical power of the method with the help of synthetic datasets where the ground truth genetic architecture is known. We used the genotypes of the widely used Drosophila Melanogaster Genetic Reference Panel (DGRP^21^, see Methods) to design our simulation scenarios. The genotype data in our scenarios comprised 1,000 SNPs with a minor allele frequency (MAF) greater than 0.1 subsampled from 2,456,752 genome-wide SNPs of the DGRP genome dataset. We considered three simulation scenarios where all are additive models containing four SNPs with marginal effects and 2 to 4 SNPs with interaction effects (see Figure 2, to better understand the overall schematic of how the simulations are conducted). The scenarios differ only in the number of interaction pairs/sets and number of SNPs involved in these interactions. Scenario 1 considers the simplest scenario where there exists only 1 two-SNP interaction in the linear additive model (called S1), Scenario 2 comprises 2 two-SNP interactions (S2) and Scenario 3 comprises 1 three-SNP interaction (S3). Note that S2 is so designed such that the set of SNPs involved in the first binary interactions has a lower marginal effect (5 and 6 in S2, see Figure 2) and SNPs involved in the second binary interactions have a higher marginal effect (7 and 8 in S2, see Figure 2). Apart from the 6 to 8 SNPs (1,2,…,8, illustrated in Figure 2) with which the model is built, we randomly add 30 or 50 other SNPs (from the set of 1000 SNPs) and conducted 100 simulations for each scenario. More details related to the simulation scenarios can be found in the Methods.

**Figure 2:**
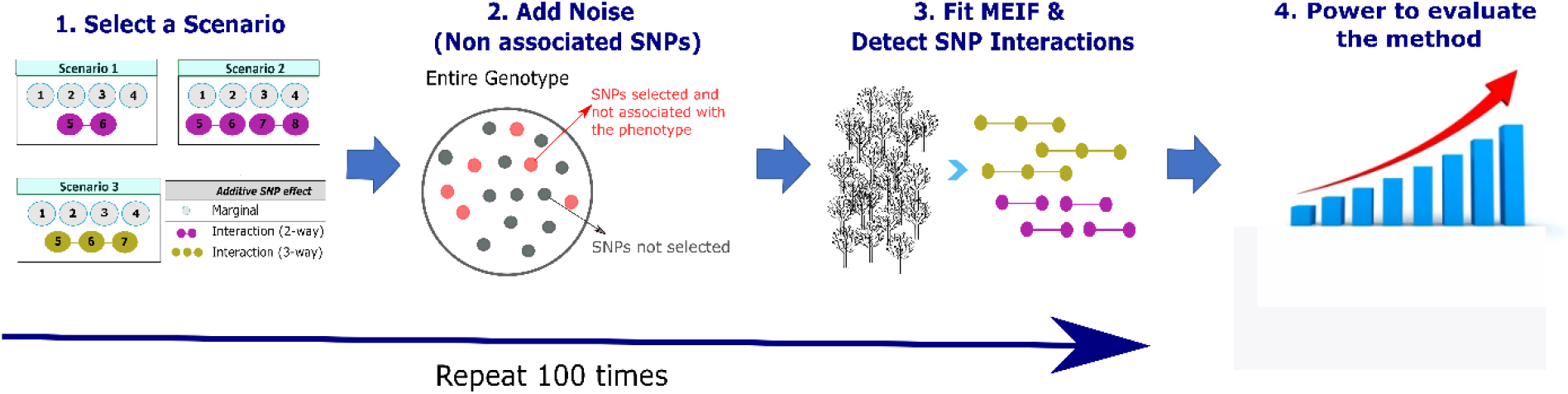
Step-by-step schematic diagram illustrating the simulation pipeline. 1) Select a simulation scenario S1, S2 or S3 involving the SNPS 1 to 8. 2) The set of causal SNPs for fitting epiMEIF in the simulated data is prepared using the SNPs with which the simulation scenario is built, and randomly selected SNPs from the set of 1000 SNPs that are not associated with the simulated phenotype (highlighted in red). These additional markers in Step 2 act as noise and help to evaluate the power of the method. 3) Fit epiMEIF (or *weighted epiMEIF*) and obtain the set of interactions. Repeat step 1 to 3, 100 times. 4) Detect the proportion of times epiMEIF can capture the true interaction (e.g., Interaction between 5 and 6 in Scenario 1) amongst the 100 iterations.

We evaluated the performance of our approach based on the proportion of simulations (out of 100) where the true interactions (example: 2-way interaction 5-6 is the true interaction in S1, see Figure 2) are captured. We call it “the overall power in capturing the true epistatic interactions” and explored the overall power for both *epiMEIF* and *Weighted epiMEIF* (see Table 1). For both the adaptation, the overall power in detecting the true interactions is consistently satisfactory ranging between **90 and 100%** across the scenarios S1 and S2. The power to detect the binary interactions in S2 is higher (2 to 4% increase) with weighted *epiMEIF* compared to *epiMEIF*. Note that despite the increased complexity in identifying higher order interactions (non-binary) in S3, we attain a power of **70 to 90%**.

**Table 1:** Proportion of simulations (out of 100) capturing the true interactions for the different Age 1 simulation scenarios under the two additional sample size cases (30 and 50).

|  | #Captured (%) |  |  |  |
| --- | --- | --- | --- | --- |
| #AdditionalSNPs | 30 |  | 50 |  |
| Method | MEIF | WMEIF | MEIF | WMEIF |
| S1 | 100 | 100 | 100 | 100 |
| S2 (SNP_5: SNP_6) | 93 | 95 | 90 | 94 |
| S2' (SNP_7: SNP_8) | 90 | 94 | 92 | 94 |
| S3 | 91 | 86 | 80 | 71 |

Since *epiMEIF* provides a score to each interaction depending on the strength of the interaction sets from the cforest, we also investigated how often the true interactions are captured as the top-ranking interactions (based on the score) in our simulation scenarios (see Supplementary Figure 1). We observed that the *Weighted epiMEIF* can capture the true interaction more efficiently compared to the *epiMEIF* for the cross-sectional data simulations (*Weighted epiMEIF* appeared in the top ranks 10 to 30% more often than *epiMEIF* in Supplementary Figure 1). The power gain is more prominent for the simulation results with 30 additional SNPs. It is noteworthy to mention here that though the “overall power” of *Weighted epiMEIF* (71%-86%) is lower than *epiMEIF* (80%-90%) in S3 (see Table 1), the power to detect the true-interactions in the top ranks is much higher in *Weighted epiMEIF* (45-80%) compared to *epiMEIF* (37-45%) (see Supplementary Figure 1c).

Overall, *Weighted epiMEIF* is more effective than *epiMEIF* as it gives comparatively robust results across all the scenarios and more often selects the true interactions as top-ranking.

#### 2.2.2 Analysis on Real Data: *epiMEIF* network construction for natural variations of heart period in flies

Testing *epiMEIF* on simulated data showed its power for capturing both binary and higher order epistatic interactions among potential causal SNPs. We therefore tested the method’s ability to detect statistical interactions from the GWAS data on cardiac performance traits in Drosophila. Drosophila is an ideal model to study heart development and adult cardiac function^22–24^. We recently investigated the genetic architecture of cardiac performance in young (1 week) flies from the DGRP population and gained insights as to the molecular and cellular processes affected^25^. Importantly, correlations observed between identified genes and cardiac dysfunction suggested a conserved genetic architecture of cardiac function in both flies and humans^25^. Leveraging on this dataset, we investigated how *Weighted epiMEIF* would detect epistasis interactions among variants associated to natural variations of heart period (HP), which measures the duration of a complete cardiac contraction/relaxation cycle. To accommodate computational requirements of *Weighted epiMEIF*, interactions were tested on variants that show significant association with the quantitative trait from the single GWAS linear mixed model (LMM), with a nominal significance threshold of 10^−5^ (*<u>3484 SNPs</u>*, see Methods). Finally, *Weighted epiMEIF* application on HP dataset led to a dense network of *35* interacting SNPs (Figure 3 and Figure 4a). We have first illustrated the statistical interaction network obtained from *Weighted epiMEIF* (Figure 3) and then illustrated the biological significance of the corresponding network with the most connected nodes (Figure 4a). Binary and higher-order statistical interactions were combined to illustrate the interaction network obtained from the *Weighted epiMEIF* (Figure 3a) and some of the four-way and three-way interactions are highlighted in Figure 3b and 3c. We have also illustrated how different allelic combinations of SNPs involved in a selected higher order interaction are associated to significant HP variations (see Figure 3d). On the contrary, the allelic combinations of SNPs randomly selected in the network (and not participating in a detected interaction) have no effect on the phenotype (see Figure 3e).

**Figure 3:**
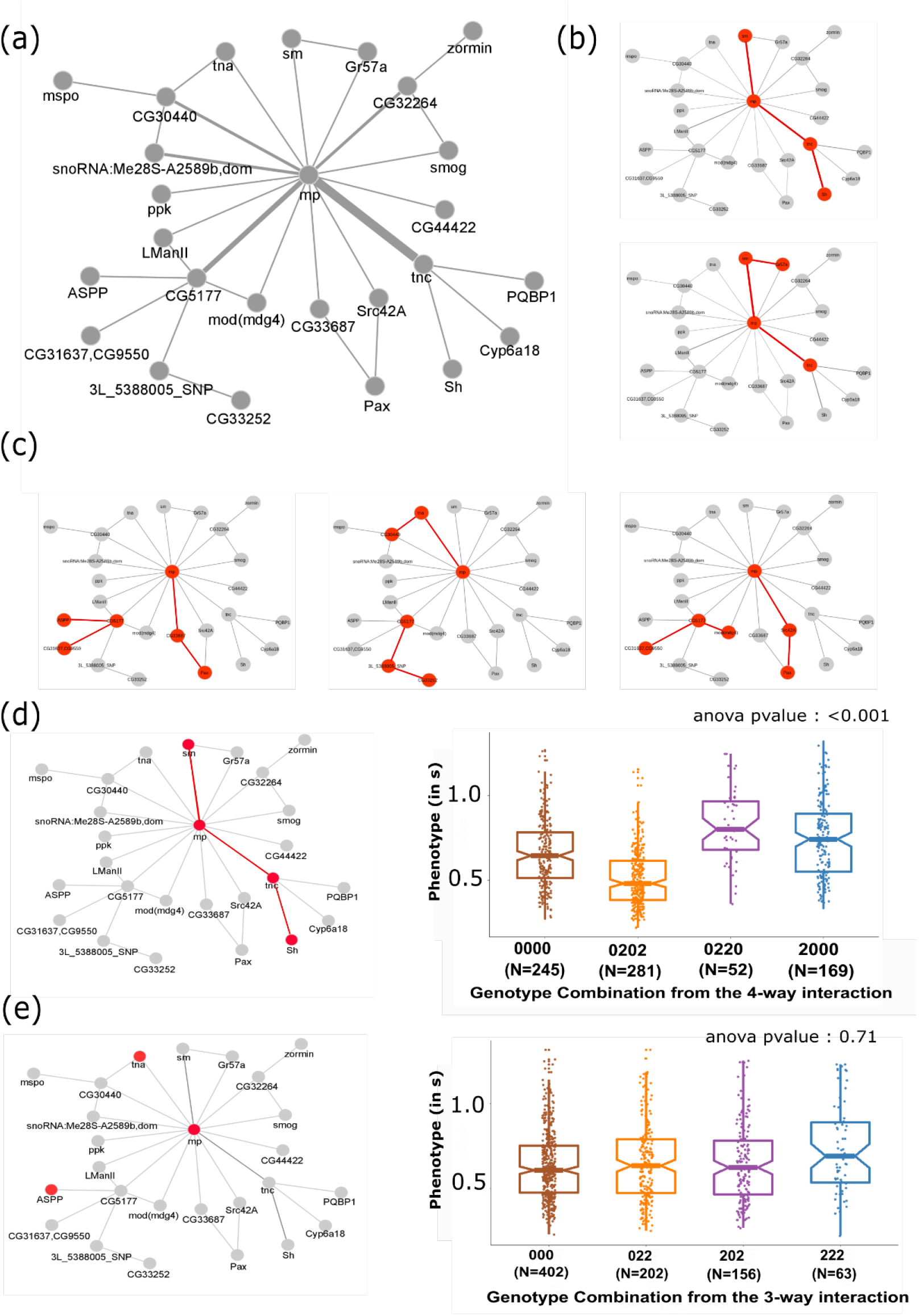
Weighted epiMEIF high-order statistical interactions detected on natural variation of heart period in flies. a) illustrates the network obtained by accumulating the detected high-level interactions obtained when weighted epiMEIF is fitted on the cardiac heart period of the DGRP population at 1 week. The different nodes denote the variants from the DGRP genotype data that are detected with the epiMEIF. Whenever the variants are mapped to a gene, the nodes are annotated with the corresponding gene name (instead of the variant name). The edge joining two nodes denotes an interaction and the thickness of the edge quantifies the number of common interactions shared by the two nodes. (b) and (c) respectively highlight (in red) some of the four-way and three-way interactions detected by epiMEIF. (d) The boxplot shows how the phenotype distribution varies across the different genotype combination arising from the selected 4-way interaction: sm-mp-tnc-Sh (in red). Max T test/anova test are performed to validate the selected high-order interaction obtained from MEIF. The p-value for the selected interaction in both tests is <0.01. (e) shows how the three-way interaction generated from a group of SNPs selected randomly from the epiMEIF network (not sharing edges/interactions) does not have any effect on the variation of the phenotype. The p-value from the anova test is 0.71, reporting an insignificant effect of the genotype on the phenotype.

**Figure 4:**
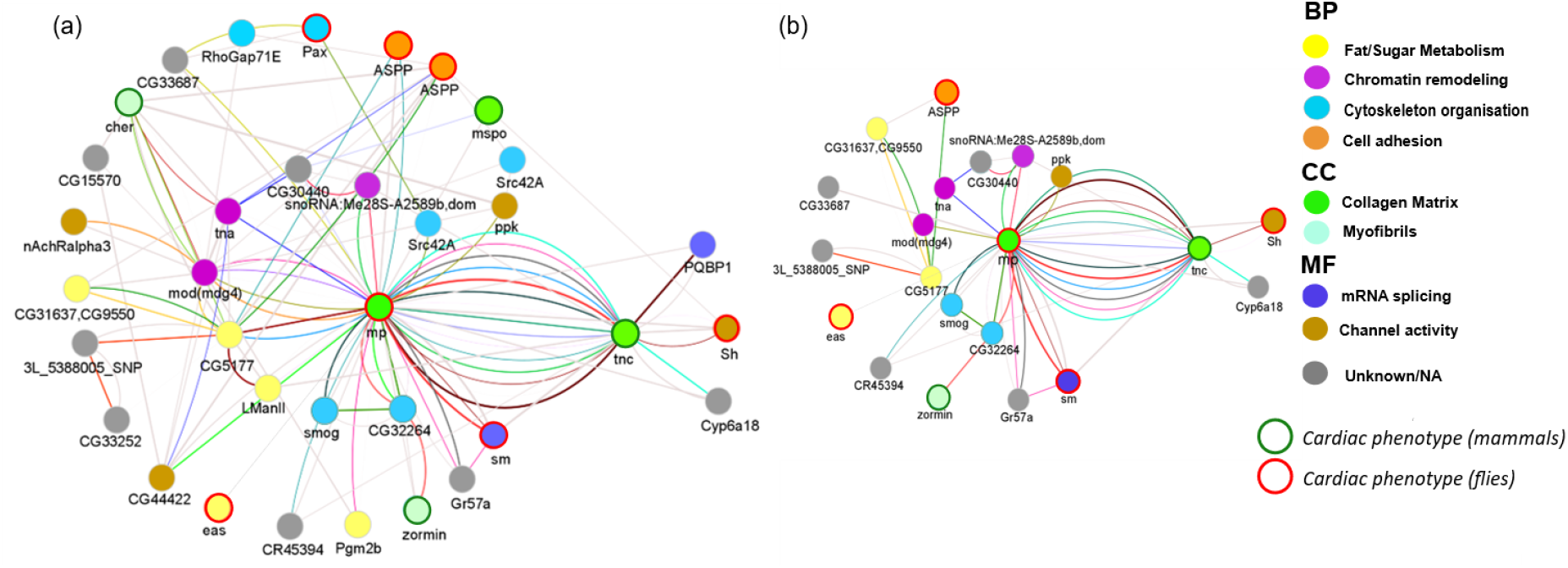
Weighted epiMEIF high-order statistical interactions detected on natural variation of heart period in flies with biological annotations. (a) shows the network obtained when weighted epiMEIF is fitted on the cardiac heart period of the DGRP population at 1 week. The different nodes denote the different variants in the weighted epiMEIF network, and wherever possible, they are annotated based on the genes to which the variants are mapped. The nodes are coloured according to their cellular and molecular functions, and the colour legend on the right denote the different cellular and molecular processes (**BP:** Biological processed, **CC:** Cellular component, **MF**: Molecular function) expressed in the epiMEIF network. The coloured boundary of the node denotes if the annotated genes have mammal orthologs associated with cardiac phenotype. (b) Part of the network in (a) that can be validated with the validation cohort using anova/Max T-test.

**Figure 5:**
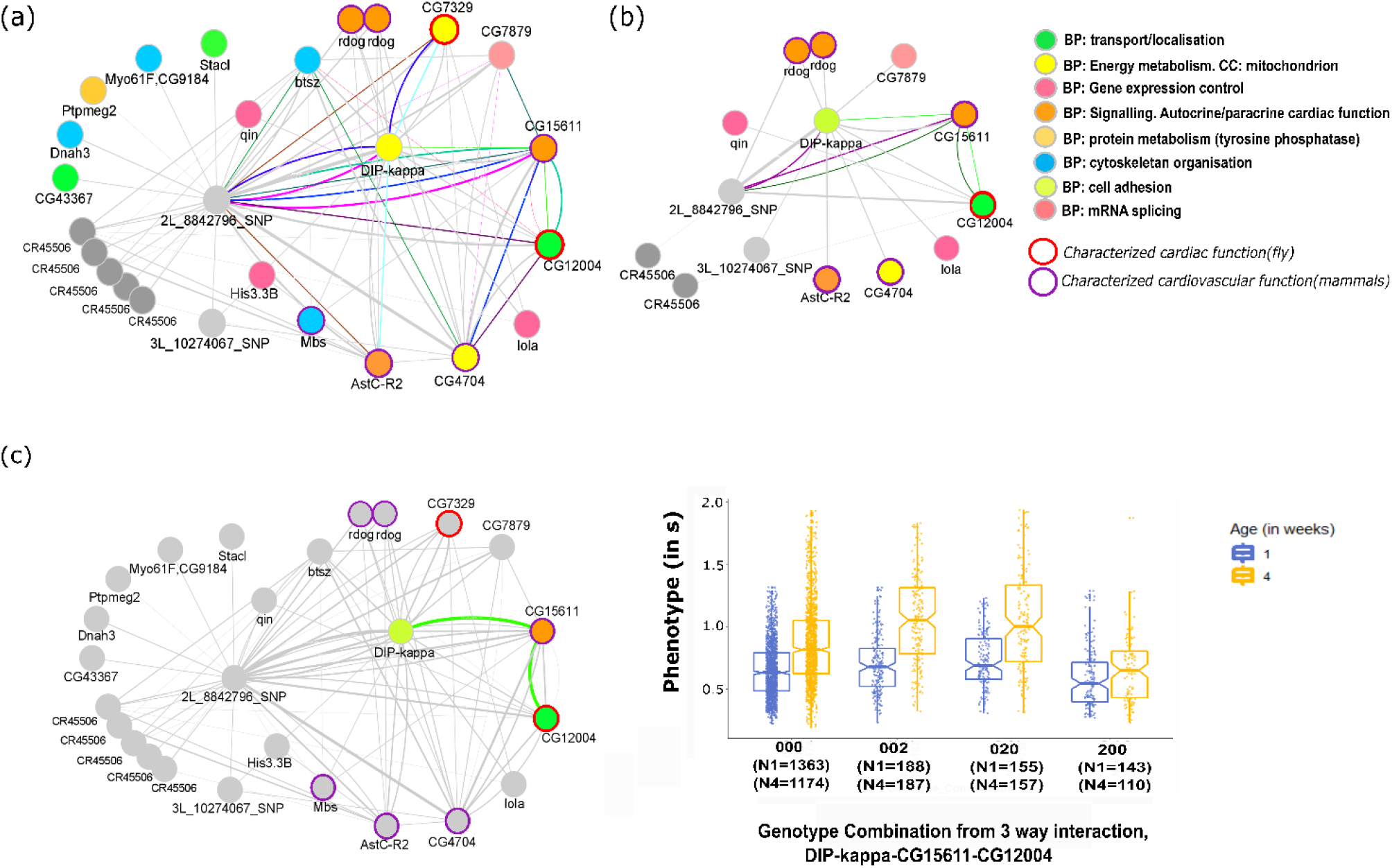
epiMEIF high-order statistical interactions detected on natural variation of heart period aging in flies. (a) network obtained when epiMEIF is fitted on the cardiac heartperiod aging data of the DGRP population (flies at 1 and 4 week). The different nodes denote the different variants in the epiMEIF network, annotated based on the genes to which the variants are mapped. The nodes are coloured according to their cellular and molecular functions. The colour legend on the right denotes the different cellular and molecular processes (BP: Biological processes, CC: Cellular component). The coloured boundary of the node denotes if the annotated genes have mammal orthologs associated with cardiac phenotype. (b) Part of the epiMEIF network in (a) that can be validated with the validation cohort using anova/Max T-test. (c) A 3-way interaction between DIP-kappa-CG15611-CG12004 is highlighted here. The boxplot distribution of the phenotype at 1 and 4 weeks against the different genotype combinations shows the aging effect of the 3-way interaction on the cardiac phenotype. The p-value from the Max-T test is <0.001.

Remarkably, a large number of genes in the network have known cardiac functions, in flies and/or in mammals, and the network connects variants in genes whose function and/or subcellular localization are highly correlated (Figure 4a, Sup. table 2). In particular, it includes several genes encoding proteins known to interact with actin - either sarcomeric (*cheerio* (*cher*, human orthologue *FLNC*)- denoted by *cher/FLNC* henceforth-, *Zormin/MYPN*) or non sarcomeric (*Src oncogene at 42A (Src42A)/FRK, cher/FLNC, CG32264/PHACTR1-2, Paxillin(Pax)/PXN, Rho GTPase activating protein at 71E (RhoGAP71E)/ARHGAP20*) – and with myosin (*smog/GPR158*). In tight interaction with the extracellular matrix (ECM), cytoskeletal proteins play a central role in the mechanical and signalling properties of the cardiomyocytes^26^. From this perspective it is remarkable that two genes encoding ECM components —*multiplexin (mp)/Col15-18* and *tenectin (tnc)/* AKAP12—have a central place in the network and share many interactions. A third collagen containing ECM constituent, *mspondin (mspo)/SPON2*, also participates in the network. Importantly, *mspo*—whose mammalian orthologue has a cardioprotective activity^27^—, is known to interact genetically with *Pax. Pax* encodes an adaptor protein which couples integrins to the actin cytoskelon^28^. *epiMEIF* interactions may thus point to a central role of the collagen-containing ECM in the mechanisms leading to variability of HP phenotype, probably by impacting on cytoskeleton dynamics. In addition, the activity of the non-receptor tyrosine kinase *Src42A*, which is involved in several cellular processes including cell adhesion and cytoskeleton organization, was shown to be regulated by *Ankyrin-repeat, SH3-domain, and Proline-rich-region containing Protein* (*ASPP*)/*PPP1R13B*^28^. Of note, several SNPs in both *ASPP* and *Src42A* are present in the network. Moreover, the network highlights other important features of natural variations of cardiac function, such as variants in *easily shocked (eas)/ ETNK1*, an ethanolamine kinase with essential cardiac function^29^, in *Shaker* (*Sh*)/*KCNA1*, a voltage gated potassium channel encoding gene involved in setting the cardiac rhythm^29^. Finally, it is worth noting that variants in *tnc* and *mp* interact with variants within *smooth* (*sm*)/*HNRNPL* and *Poly-glutamine tract binding protein 1 (PQBP1*), two genes involved in mRNA splicing, suggesting that regulation of ECM components through mRNA splicing impinges on natural variation of HP.

##### Validation of the results

We then tested if the *epiMEIF* network for heart period was replicated in an independent cohort. Twenty DGRP lines —not included in the first dataset and therefore not used in building the statistical interaction network — were analysed for cardiac phenotypes at 1 week. We used Max-T and anova tests to test whether *epiMEIF* interactions were replicated in this independent population. Note that the interaction network comprised 45 binary interactions that are not part of any higher order interactions, 33 three-way interactions that are not part of any four-way interactions and 2 four-way interactions (see Supplementary Figure 2a and Supplementary Table 2). For each interaction set of size n, anova/regression-based test detects if the n-snp interaction is statistically significant in the validation data (see Methods for more details). Max-T test, on the other hand, tests if the SNP combinations in each high-order interaction (size n) give rise to a significant impact on variation of the phenotype in the validation data and if the impact of this SNP combination is better than any random combination of n SNPs (see Methods for more details). We retained those interactions which performed well either with the anova test or Max-T test. Note that approximately 18% (8/45) 2-way and 5% (2/33) 3-way interactions could not be tested with the validation cohort due to lack of observations. Amongst the one tested 14% (5/37) 2-way interactions, 42% (13/31) 3-way interactions and 100% (2/2) 4-way interactions could be validated (see Supplementary Figure 2b and Supplementary Table 2). The biological significance of the part of the *epiMEIF* network that can be validated with the validation cohort is represented in Figure 4b. Note that only a part of the network could be validated because the reduced number of lines in the validation cohort made it less powered to test all the interactions. Nevertheless, the main characteristics of the network (in Figure 4a) are found reproduced in this independent population, including the centrality of *multiplexin (mp*) and its interactions with *tenectin* (*tnc*) and *Shaker (Sh*) (see Figure 4b). More details on the interactions in the network can be found in Supplementary Table 3.

### 2.3 Application to longitudinal data

We have shown the efficacy of our approach with cross-sectional datasets. Noticeably, the *epiMEIF* method can be extended for detecting genetic interactions from longitudinal datasets (see Methods). We have rarely encountered genetic interactions being tested on longitudinal datasets in literature. Malzahn et al.,^30^ tested for gene-gene interaction using longitudinal nonparametric association test on Framingham Heart Study (FHS) cohorts, but they did not test for higher-order interactions. Our approach is particularly interesting because it also presents a way to explore higher-order interactions in longitudinal data. Similar to cross-section data above, we have tested the efficacy of our approach in longitudinal datasets *via* simulations and real data applications.

#### 2.3.1 Analysis on Simulated Data

We evaluated the statistical power of the method on longitudinal datasets using simulation scenarios similar to those used with cross-section data. Here as well, we have 3 simulation scenarios - AS1, AS2 and AS3. While AS1 considers the simplest scenario with 1 binary interaction, AS2 considers a comparatively complicated scenario with 2 binary SNP interactions. AS2 is designed as earlier; one set of SNPs involved in the interaction has a lower marginal effect and the other set of SNPs has a higher marginal effect. AS3 comprises 1 three-SNP interaction (see Methods for more details). We have conducted our simulations 100 times for each scenario, and we run each scenario with 30 or 50 additional variants, randomly chosen from the genome data (with 1000 SNPs), same as earlier. Note that we have conducted the longitudinal simulations only with *epiMEIF*. *Weighted epiMEIF* cannot be easily adapted for longitudinal dataset as deciding a weight for the “Time” covariate (that captures the “longitudinal” component of the data) is challenging. Assigning a heavy weight to “Time” can lead to trees where “Time” is often selected at the root node, whereas assigning a low weight can end up with trees where “Time” is never selected for any node (see Methods for more details). Hence, to avoid complications we focus on *epiMEIF* application in longitudinal datasets.

As in the previous simulations, we evaluated the performance of *epiMEIF* on longitudinal data based on “the overall power in capturing the true epistatic interactions” (see Table 2) and power to capture the true interactions as top-ranking (see Supplementary Figure 3). The “overall power” is quite high (**95 to 98%)** across all the scenarios (see Table 2). Both the “overall power” and the “power to capture the true interactions in the top ranks” are similar across the two situations-with 30 and 50 additional variants (see Table 2, Supplementary Figure 3). However, *epiMEIF* is less efficient in capturing the interactions involving SNPs with lower marginal effect (power to capture the true interactions in top ranks is as low as **10 to 20%** for the interaction pair 1-2 in scenario AS2 in Supplementary Figure 3). Having acknowledged that, the overall power to capture true interactions is quite satisfactory across all the scenarios and exploring higher order interactions with synthetic longitudinal data presented here is novel in GWAS.

**Table 2:** Proportion of simulations (out of 100) capturing the true interactions for the different Aging simulation scenarios under the two additional sample size cases (30 and 50).

|  | #Captured (%) |  |
| --- | --- | --- |
| #AdditionalSNPs | 30 | 50 |
| Method | MEIF | MEIF |
| AS 1 | 97 | 97 |
| AS 2 (SNP_1:SNP_2) | 98 | 98 |
| AS 2 ( SNP_5:SNP_6) | 97 | 98 |
| AS 3 | 96 | 94 |

#### 2.3.2 Analysis on Real Data: *epiMEIF* network construction for natural variations of heart period aging in flies

To further evaluate our method for identifying epistatic interactions from longitudinal data, we analyzed the aging of the heart function in the DGRP population. Several cardiac aging studies in Drosophila have revealed striking similarities with mammals, both in terms of heart physiology and transcriptional changes^22,23,31,32^. This highlights the conserved nature of cardiac aging across organisms. We analysed 165 of the DGRP fly lines included in the previous dataset for heart period at 4 weeks of age (12 individuals per line, see Methods). Overall, there is a marked increase of heart period in the DGRP from 1 week to 4 weeks, in agreement with previous observation (see Supplementary Figure 4). Both 1 week and 4 weeks data on the 165 DGRP lines were used to identify epistatic interactions between variants associated to natural variations of heart period during aging. Similar to the previous 1-week study, *epiMEIF* interactions were identified on variants that show significant association with the aging of HP from the single GWAS longitudinal LMM, with a nominal significance threshold of 10^−5^ (1682 *SNPs*). Finally, *epiMEIF* application on HP aging dataset led to a dense network of *26* interacting SNPs; comprising 47 two-way that are not part of any 3-way interactions and 12 3-way interactions (Figure 4a, Supplementary Table 4). Strikingly, the *epiMEIF* network comprises tightly interconnected variants within genes involved in diverse biological processes, many of which have characterized cardiac function, either in flies or in mammals. In particular, the network includes variants in 3 genes encoding signalling pathway components, namely *Allatostatin C receptor 2 (Astc-R2*), the somatostatine receptor orthologue; *red dog mine* (*Rdog*), an ATPase-coupled transmembrane transporter orthologous to *ABCC4;* and *CG15611*, the Rho guanine nucleotide exchange factor *ARHGEF25* orthologue. Noticeably, *rdog* and *CG15611* orthologues have known autocrine or paracrine cardiac functions either in human or mice ^33–35^. This suggests a primary role of natural variations of cardiac signalling properties in heart senescence. In addition, the numerous interactions that these SNPs engage in within the network suggest their involvement in relation to the other variants associated with the aging of cardiac function. Among these, one SNP into *Dpr-interactingprotein κ (DIP-kappa*) – encoding a cell adhesion protein orthologous to *LSAMP-* interacts with 13 variants into 12 genes. One SNP into *CG4704*, which encodes the fly orthologue of the human *MCU1* (Mitochondrial Calcium Uptake 1) - interacts with 10 variants into 9 genes. *MCU1* is a regulatory subunit of the Mitochondrial Calcium Uniporter which plays a central role in calcium import into the mitochondrion and in mitochondrial calcium ion homeostasis^36^. This suggests a central function of these components in heart period aging. The most connected node corresponds to a SNP that is more than 10kb away from any gene, precluding its annotation. The network additionally includes SNPs in genes involved in cytoskeleton organisation (*Myosin binding subunit (Mbs), bitesize (btsz)*) in carbohydrate transport (*CG12004/TMEM184*)), in energy metabolism (*CG7329/LIPM*) and in mRNA splicing (*CG7879/RBM12*). Strikingly, 5 variants within 150 bp upstream of the lncRNA *CR45506* are retrieved in the network, suggesting a major involvement for this lncRNA in the process, throughout interactions with several other members of the network. Taken together, these SNPs and their interactions allow us to draw some characteristics of the genetic architecture of natural variations in the aging of cardiac function. More details on the interactions in the network can be found in Supplementary Table 5.

##### Validation of the results

The twenty DGRP lines not included in the first dataset were also analyzed at 4 weeks of age, thus providing a validation set for cardiac aging which was used to test for replication of the interaction using Max-T/ Anova test in this independent cohort. 75% (35/47) two-way interactions and 58% (7/12) three-way interactions could be analysed in this validation cohort (see Supplementary Table 6). Of them, 43% two-way (15/35) and 43% three-way (3/7) were replicated (Figure 4b). Several features of the network were replicated in the independent cohort, including the involvement of *rdog, CG15611 and Astc-R2*. This also confirmed the central positioning of *DIP-kappa*, whose implication in natural variations of cardiac aging therefore warrants further investigations.

## Discussion

Here, a new method is proposed for epistasis detection in large scale association studies with complex genetic traits using mixed effect conditional inference forest (*epiMEIF*). This method captures higher-order SNP interactions based on the tree structure in the cforest, and combined with mixed models, can handle a wide range of complex GWA studies. The effectiveness of *epiMEIF* is verified in extensive simulation scenarios reflecting a wide spectrum of complex models and with real datasets, illustrating its power for epistasis detection from both cross sectional and longitudinal data. The additional testing strategies applied a posteriori to the conditional inference forest in *epiMEIF* not only safeguards the method from detecting false positive interactions but also increases the reliability of the final selected interactions. Moreover, by demonstrating the ability of the approach to replicate in an independent cohort we ensure that our approach does not suffer from winner’s curse^37^ which is quite common in GWAS. The ability of the approach to validate part of the higher-order interactions in an independent cohort also supports its’ competency in detecting higher-order interactions. Additionally, for the cross-sectional datasets, we proposed an adaptation of *epiMEIF*, named *Weighted epiMEIF* that allows identifying genes associated with weak marginal effect variants. This is an important addition to the ‘traditional’ epistasis approaches that are biased towards variants with strong marginal effects^7,8,20^. Similar to LMM, a major advantage of *epiMEIF* is their ability to effectively account for unwanted correlation between samples, thereby correcting for confounding factors such as population structure^38^ or hidden covariates^39^ and making it easily applicable for human GWAS studies as well. However, unlike the standard LMM, *epiMEIF* can jointly model genetic effects of multiple loci or markers on the readout. This is particularly important because recent works have revealed that often, the single-locus association models are insufficient to explain the heritable component of complex traits^40,41^. The proposed approach has proven to overcome the caveats of existing GWAS approaches and though we have shown its applicability for identifying associations and epistasis detection, it can be also used for prediction^42,43^ and feature selection^20,44^. We however acknowledge that the method is not scalable to the entire genome dataset and do not provide the exhaustive list of interactions from the entire list of variants. Interestingly, the *epiMEIF* generate dense genetic interaction networks that are creating hubs around some focal genes. Part of this distinctive topology may be attributable to the nature of tree formation in the cforest algorithm. It is difficult to determine if this topology is also due to the nature of the genetic interactions that may underlie this statistical network. The analysis of the networks obtained from the study of natural variation in cardiac function undoubtedly sheds light and should indicate that the statistical interactions reveal an underlying biological functionality. Indeed, the vast majority of the interactions identified affect genes whose products are involved in the cytoskeleton (sarcomeric and non-sarcomeric) and in its dynamics and, in interaction with the extracellular matrix, must participate in the mechanical and signalling properties of cardiomyocytes.

In systems biology there is an emerging interest in understanding the genetic mechanisms underlying the study of longitudinally measured phenotypes^45,46^. Additionally, it has been often discovered that complex diseases are the results of interaction between a large number of units, and therefore, are more likely to be associated with genes that are well connected in the network^47^. epiMEIF can comprehensively model the dynamics of longitudinal data and crosssectional data and offer to reveal the relationships between multiple variants, revealing the extensive networks of genetic interactions that are causing the complex diseases. The possibilities offered by *epiMEIF* will allow approaching questions that were previously difficult to address, as tools for the identification of epistatic interactions in complex datasets were lacking. In particular, we do not know when biomolecular interactions actually produce patterns of statistical epistasis, nor do we know how to biologically interpret statistical evidence of epistasis, since statistical interaction likely does not automatically entail interaction at a biological or mechanistic level. Progress towards these important questions will provide a framework for using genetic information to improve our ability to diagnose, prevent and treat common human diseases^48^. Hence, our approach provides the foundation for extracting higher-ordering statistical interactions flexibly from any type of dataset, that will guide the biologists in formulating their hypothesis. Eventually, network analysis tools^49^ may be useful to biologically interpret the statistical evidence of epistasis and bridge the gap between statistical and biological epistasis network.

## METHODS

We propose an approach for epistasis detection using mixed effect conditional inference forest (*epiMEIF*) to identify the genetic variants and their epistatic interactions responsible for the complex quantitative traits. The novelty of this approach lies not only on the epistasis detection but also on the amalgamation of mixed effects modelling and conditional inference forest (cforest). We have divided the explanation of the *epiMEIF* method into four parts: 1) we explain the *MEIF* model, 2) we illustrate how the tree structure in the conditional inference forest (cforest) is utilized to detect high-order SNP interactions that impact the variation of the phenotype, 3) we explain how MEIF can be adapted to weighted MEIF, this is particularly useful for cross-sectional dataset, and 4) we show how the interactions from the MEIF can be validated using independent statistical tests thereby leading to a final Interaction Score Matrix (*F*).

### 1. Mixed Effect Inference Forest (MEIF)

Our method is primarily inspired by the Mixed effect random forest (MERF) proposed by Hajjem et al^50^. It is an ensemble of mixed effects model and random forest where the application of random forest is extended to clustered data for a continuous outcome. The following model is proposed in MERF:

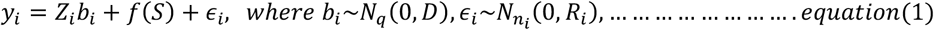

In the equation above:

- *i* is the cluster index. Assume that there are *K* clusters in the training data (e.g., in the DGRP dataset, cluster denotes the DGRP strain).
- *y_i_* is the target variable. For cluster *i*, there are *n_i_* observations and *y_i_* is the *n_i_* × 1 vector of responses (e.g., in the DGRP dataset, *y_i_* is the phenotype of interest).
- *S* is the set of covariates/markers and it is assumed to be *N* dimensional, e.g., in the DGRP data the *N* features are the causal SNPs in the genotype data.
- The non-linear function *f* is learned using a random forest.
- *Z_i_* is the random effect features and it is assumed to be *q* dimensional,(i.e., there are *q* features). Hence, *Z_i_* = [*Z*_*i*1_, *Z*_*i*2_,…, *Z_in_i__*]^*T*^ is the *n_i_* × *q* matrix of random-effects covariates for cluster *i*.
- *b_i_* is the random effect coefficient for cluster *i*.
- *ϵ_i_* is independent, identically distributed (*iid*) noise.

We adapted the above model (*equation*(1)) in our method such that it can also include some fixed-effect covariates (*X_ij_*) that is not part of the random forest or random effects (see ∑*a_j_X_ij_* in *equation*(2)) below) and that have ideally linear additive effects on the response. Moreover, our aim was to implement the current model on genomic data where some of the variants are not properly sequenced for all the samples and suffer from missing data issues. To fit a random forest on genotype data with missing values one usually need to impute the genotype data and then apply the above method. To avoid that, we applied conditional inference forest(cforest)^51^ instead of traditional random forest (see *MEIF model* in *equation*(2)) and hence, call our method mixed effects conditional inference forests (*MEIF*). Cforests, developed by Torsten Hothorn et al. in 2006^51^, can be considered as an alternative version of the original random forests where the ensemble algorithms uses conditional inference trees as base learners. Similar to traditional random forests, Cforest also constructs a collection of classification/prediction trees and aggregates them into one robust classifier/prediction tool. The deterministic tree-growing algorithm in these methods rely on two vital stochastic rules: (i) each tree is trained on a bootstrap sample of the original sample; (ii) at each node in the trees, the best split is chosen from among a randomly selected subset of the predictor variables instead of the full set^8^. However, traditional random forests are based on CART (Classification and Regression Tree) that has been criticized for its bias in variable selection and often suffer from overfitting and a selection bias towards covariates with many possible splits or missing values^52^. Cforest based on conditional inference trees select variables in an unbiased way and the partitions induced by this recursive partitioning algorithm are not affected by overfitting and surrogate splits are exploited when covariates have missing data. More details on Cforest can be found in Hothorn et al,.^51^

The *MEIF* model used in our method can be defined as follows:

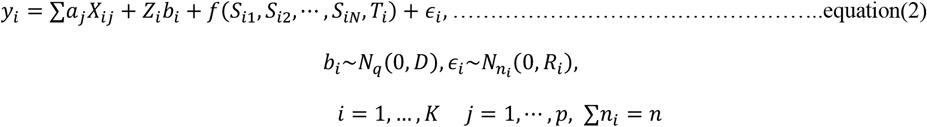

where

- *y_i_* = [*y*_*i*1_, *y*_*i*2_,…, *y_in_i__*]^*T*^ is the *n_i_* × 1 vector of responses for the *n_i_* observations in cluster *i*,
- *X_ij_* is the *j^th^* fixed-effect covariate for cluster *i*. 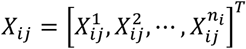 is the *n_i_* × 1 vector of *j^th^* fixed-effect covariate values for cluster *i*, and ***X_j_*** = [*X*_1*j*_, *X*_2*j*_,…, *X_k_j__*]^*T*^ is the *n* × 1 vector of *j^th^* fixed-effect covariate values over all clusters (e.g., in the DGRP dataset, *X_ij_* comprises the variants like the date on which fly is dissected).
- *Z_i_* is the *n_i_* × *q* matrix of random-effects covariates that we do not intend to include in the cforest.
- *S*_1_, *S*_2_,… *S_N_* are the marker variables in our study and *T* capture the time or the longitudinal component of the data (*T_i_* denotes the time at which *y_i_* is observed). *f*(*S*_*i*1_, *S*_*i*2_,…, *S_iN_, T_i_*) is ideally estimated using conditional inference forest (cforest).
- *a_j_* is fixed effect of the *j^th^* covariate,
- *b_i_* = [*b*_*i*1_, *b*_*i*2_,…, *b_iq_*]^*T*^ is the *q* × 1 unknown vector of random effects for cluster i,
- *ϵ_i_* = [*ϵ*_*i*1_, *ϵ*_*i*2_,…, *ϵ_in_i__*]^*T*^ is the *n_i_* × 1 vector of error variables.

Note that we incorporate the cforest on the markers and time covariate only as we would like to capture the interactions of these variables. The covariate time (*T*) can be omitted for cross-sectional data.

Pooling data across all the clusters, we have the following *MEIF* model for cross-sectional dataset:

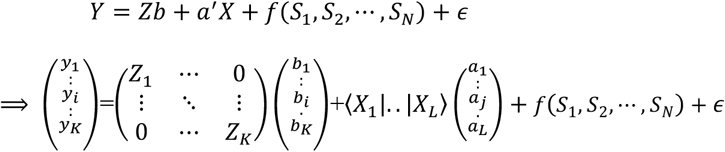

For longitudinal dataset, the above models can be written as follows:

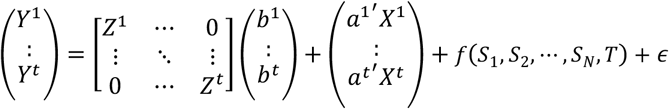

where, *t* denote total number of time points in the longitudinal data and superscript t for each component denote the corresponding response and predictors at time point t. Note that, similar to the original article^50^ an expectation-maximization (EM) technique is used to fit the *MEIF* model.

### 2. Epistatic interactions detection with Epi-MEIF

The random forest technique is an ensemble learning technique for conducting classification and prediction analyses^53^. Apart from functioning as a prediction or a classification tool, the random forest also acts as feature selection tool. It provides importance score corresponding to each variable used in the model that measures their contribution to the predictive accuracy^8^. In the context of genetic association studies, this importance score can be used to discover variables, i.e., SNPs, that are most predictive of the disease status (thereby likely to affect disease susceptibility) or the SNPs that are most predictive of the phenotypic variability. Unlike the single GWAS model, the phenotype predicted using random forest (RF) accounts for the combined/interaction effect of various SNPs^8^. In this regard, another unique feature of RF is that the tree structure in the RF can be exploited to detect the SNP interactions. The nature of the trees constructed allows for interaction in the sense that each path through a tree corresponds to a particular combination of values taken by certain predictor variables, thus including potential interactions between them. If a particular path (or a sequence of covariates) occurs more often than the others in multiple trees of the random forest, we can claim the group of variables in the path are in interactions. Note that the method is not scalable to a genome-wide setting. Hence, to retrieve the set of potential causal SNP inputs for epiMEIF, we propose to apply a single-locus association (using LMM) that first filters the SNP markers that have a significant effect on the phenotype. This can help in alleviating the computational burden of epiMEIF (illustrated in Figure 1).

#### Detecting epistasis in Cross-sectional Data

Supplementary Figure 5 illustrates the mechanism of epistasis set construction from the random forest/conditional inference forest in the cross-sectional dataset^54^. Precisely, if two SNPs, C and E have a large epistatic effect then C and E combination will appear more often in the same branch of a tree than in other branches or trees (see Supplementary Figure 5). This combination will thus form a parent/descendant (child, grandchild, and so on) pair. On the contrary, if two SNPs, like B and C, have large but independent main effects on the response variable, they will also appear frequently within the same tree, but in different branches (that are not descendant pairs) (see Supplementary Figure 5). Thus, the descendant pairs that occur most frequently in the RF can be recognized as possible pairwise epistatic interactions. Similarly, C, D and E are expected to occur more often in the same branch of the tree if they have large epistasis effect (see Supplementary Figure 5). The tree structure thus allows to identify higher-order (more than pairwise) interactions. Supplementary Box 1 further elaborates the different steps of the MEIF.

##### Supplementary Box 1: MEIF Algorithm

Step 1: Fit the MEIF as shown in equation(2). Step 2: Extract the cforest component 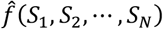 from the fitted MEIF model, where in each forest the decision is taken based on the cumulative decision from n trees. Following the mechanism in Supplementary Figure 5 each tree gives rise to a potential interaction set {*Q_t_, t* ∈ {1, 2, … *n*}}. Compiling all the interactions sets {*Q_t_, t* ∈ {1, 2, … *n*}}, SNP Interaction Matrix (*I*) is computed, that measures the strength of each interaction set based on their frequency of occurrence in the cforests. Step 3: Since cforest is a bagging algorithm where the individual trees are built from bootstrap samples, different iteration of cforest may give rise to different SNP Interaction Matrix (*I*). Hence, to enhance the stability of the final interactions sets, we propose to fit the MEIF/cforest 10 times and then obtained the interaction sets that are occurring with high scores in 90% of the forests (9/10 forest). Step 4: The interaction sets along with their pooled interaction score from the 10 forests are utilized to construct statistical epistatic clusters (see Figure1 and Supplementary Figure 7).

The mechanism illustrated in Supplementary Figure 5 will finally yield the list of all potential interaction sets from the forest. We compute a SNP Interaction metric (*I*) that measures the strength of each interaction set based on their frequency of occurrence in the same branch of the forest (see Supplementary Box 1, for more details). Note that the above mechanism is relevant for identifying interactions in cross-sectional phenotype-genotype studies and it needs to be extended for longitudinal case studies.

#### Detecting epistasis in Longitudinal Dataset

Ideally, the longitudinal aspect of the dataset is captured with the variable “Time”. Since our primary objective is to capture the epistatic interactions that are responsible for the change of the phenotype over “Time”, we focus on those branches of the trees that have “Time” in at least one of the child nodes (see Supplementary Figure 6). If A, C and D appear more often in the same branch of a tree as ancestor of “Time”, then A, C and D are expected to have epistatic effect having an impact on the longitudinal variation of the phenotype (see Supplementary Figure 6). Each path of SNPs from root to leaf preceding the node having the “Time” covariate is a possible interaction. Certain combination of the SNPs that appear more frequently in the same branches are believed to interact with each other. Adopting this intuition, we build SNP Interaction metric (*I*) that measures the strength of each interaction set based on their frequency of occurrence in the same branch with “Time” in the forest (see Supplementary Figure 7).

Note that we are essentially interested in those interactions where the temporal variation of the phenotype changes significantly from one genotype combination to another. Supplementary Figure 7a shows the phenotype distribution of a group of individuals across the different genotype combinations arising from a two-SNP interaction in the Drosophila Cardiac Genetics dataset (see details on the dataset below) observed at two time points. In this article, 0 and 2 denote the major allele and minor allele, respectively. The highlighted combination 00 denote the major allele combinations of the binary interaction. We consider the population belonging to the major allele combinations (example: 00 in Supplementary Figure 7a) as the reference population in our dataset. We are interested in identifying the SNP interactions where the longitudinal phenotypic trend of the alternative allele combinations (example: 02, 20, 22 in Supplementary Figure7a) is significantly different from the major allele combinations. To address that, we remove the longitudinal trend of the reference population from the entire dataset and then explore the epistatic interactions using MEIF. This is illustrated in Supplementary Figure 7a. Supplementary Figure 7b shows the step-by-step mechanism to compute the interactions sets and their strength of interaction from the forest in a longitudinal data. The different steps (5 steps) in Supplementary Figure 7b can be summarized as follows:

1. An example of a tree of interest in MEIF is shown here (where Time is not in the root node). SNP A, SNP C and SNP D forms a tree with Time as the child node because the longitudinal phenotype trend of genotype combination 020 (SNP A=0, SNP C=2 and SNP D=0) is significantly different from 000 (SNP A=0, SNP C=0 and SNP D=0). 2) and 3) Whenever the SNPs like SNP A, SNP C and SNP D appears in combination with Time (multiple times), one can say that they are in epistasis, and they are attributing to the longitudinal variation of the phenotype. All possible interactions with Time from the *i^th^* tree are listed here. The proportion of times a set of SNPs appear together in one tree is captured in the SNP Interaction matrix (*I_i_*) illustrated here. 4) The SNP:SNP (*I*) Interaction matrix captures the strength of interaction between all the SNPs from all the trees in the forest, and it is utilized to build epistasis clusters as shown in 5). The Time: SNP (*A*) interaction matrix captures the strength of interaction of SNPs with Time, thereby identifying the vital SNPs that affect the longitudinal variation of the phenotype. The SNP Interaction metric (*I*) can help us build the epistasis clusters/network of genetic variants as illustrated in part 4 of Supplementary Figure 7b.

As illustrated in Supplementary Box 1, we have fitted 10 cforests or MEIFs and then obtained the interaction sets that are occurring with high scores in 90% of the forests (9/10 forest). Though bagging^53^ in RFs, that is, learning individual trees on bootstrap samples and aggregating these over the resulting ensemble, yields the necessary stability to the *MEIF* approach, we further boost the stability of the final interaction sets by pooling across multiple cforest as proposed above.

### 3. Weighted Conditional Inference Forest

A notable shortcoming of tree-based methods is that they are dependent on marginal effects. Random forests are essentially bagging approach which begins with the bootstrap sampling of individuals from the original dataset followed by feature selection for the construction of each node of the tree. To do so, instead of considering all variables from the initial GWAS dataset, a random subset of variables is picked without replacement. Hence at the tree learning step, the algorithm looks for a single SNP that well discriminates cases from controls or different groups of phenotypes. In practice, this is equivalent to looking for SNPs with high marginal effects. To overcome this, instead of randomly drawing the variables, we inject weights to each variable used in the random forest. The weights indicate the probability of selecting the variable (or markers in our situation) during the construction of the node of the tree.

#### Weight design

The Linear Mixed Model (LMM) applied at the first stage prior to *MEIF* application provides the significance of each variable based on a single marker association test. We utilize this to rank the SNPs based on the size of their marginal effects and the p-value from the LMM. The weight assigned to each SNP is inversely proportional to the rank of the SNP (i.e., highly significant SNPs have lower weights). This ensures that SNPs with higher marginal effects are drawn less during the tree construction, thereby increasing the chance of capturing interactions involving SNPs that have low marginal effects^55^. The relationship between the weights and the p-values is illustrated in Supplementary Figure 8 where the logarithm of the p-values of 3484 SNPs selected after single GWAS on the Heart-period variations from the DGRP data is plotted. The weights assigned to each SNP is inversely proportional to the p-values and is plotted below. The rank of each SNP (based on the p-values) is shown in the x-axis.

Note that in the cforest implementation we could control the weights of sampling only for the root node. For the subsequent child nodes each covariate has the same chance of getting sampled just like the traditional random forest. One hurdle for the longitudinal data is to decide on the weight for the “Time” covariate. Assigning a heavy weight can lead to trees where “Time” is often selected at the root node, whereas assigning a low weight can end up with trees where is “Time” is never selected for any node. So, we avoided the weighted cforest or *Weighted epiMEIF* application for longitudinal data and have applied it only for cross-sectional simulated and real datasets.

### 4. Epistasis Network Validation

We propose to perform an independent statistical test on the interaction lists obtained from the *MEIF*. The objective of this testing step is to further validate the interactions obtained from *MEIF* and help us get rid of false positives, if any. Within this statistical validation framework, we intend to address 2 primary questions: 1) Do the SNP combinations in each high-order interaction (selected from the *MEIF*) give rise to a significant impact on variation of the phenotype? (Anova test) 2) Is the impact of this SNP combination better than any random combination of N SNPs, where N is the size of the high-order interaction (under investigation)? (Max-T test). To address this, we compare the interaction sets from *MEIF* with random subsets of SNPs from the whole genome and show that the interaction sets indeed have a higher impact on the variation of the phenotype compared to any random subset of SNPs. We first explain how the interactions are validated using Max-T test and then explain anova/regression-based tests can be used for validation.

#### Max-T test

The proposed strategy for statistically validating the list of interactions using Max-T test is schematically shown in Supplementary Figure 9 and Supplementary Box 2. An example of a plot showing the impact of a 3 SNP epistatic interaction on the aging of the phenotype in the cardiac aging fly dataset is presented in Supplementary Figure 9 Note that the aging trend (Change in Phenotype from Age 1 to Age 4) for the combination of all 2’s (minor allele combination) is significantly different from the aging trend for the combination of all zeros or major allele combination (highlighted by dotted blue curve). We are interested in highlighting such SNP combinations in this step. Supplementary Box 2 elaborates on the step-by-step process of validating an epistasis set of N SNPs (generated from MEIF).

##### Supplementary Box 2: Validating a set of N-way interactions using Max-T test

Step 1: Randomly draw any N SNPs from the whole genome excluding the causal variants (n times, 1000 in our case). Step 2: For each N-way SNP set compute and evaluate the Max-T test using the following:

a. *N* SNP interaction give rise to 2^*N*^ genotype combinations denoted by the set *G* = {1,…,2^*N*^}.
b. Fit the regression:

- For longitudinal dataset: (*Y_i_*(*t*) = *β_i_ t* + *α* + *ϵ*), where *i* ∈ *G*, *Y_i_* (*t*) denote the phenotype of the population having genotype *i* at time point t and *β_i_* capture the effect of the genotype *i* on the longitudinal variation of the phenotype.
- For cross-sectional dataset: (*Y_i_* = *β_i_* + *α* + *ϵ*), where *i* ∈ *G*, *Y_i_* denote the phenotype of the population having genotype *i* and *β_i_* capture the effect of the genotype *i* on the variation of the phenotype. Note that we assume that the genotype combination 1 as the baseline here denoting the combination arising from the interactions of major alleles. For instance for the DGRP data, SNP value 0 denote the major allele and 1 in *G* denote the combinations with all 0’s.
c. We test the following to asses if the effect of any alternative genotype combination is significantly different from the major allele combination:

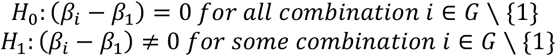 The test statististics used to evaluate the above:

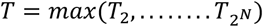

where *T_i_* is statistics computing the difference in effect of the alternative genotype from the baseline genotype on the variation of the phenotype.

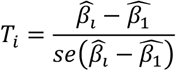 Say, *β* = {*β*_1_,…, *β*_{_2^*N*^__}} and 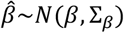, *where β* denotes the effect of all the genotype combinations on the variation of the phenotype and ∑_*β*_ is a diagonal matrix that captures the variance of these effects. The effect due to one genotype is independent of the effect of the other. Hence, we can reformulate:

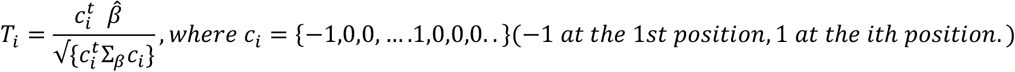 Say, 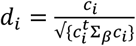, hence

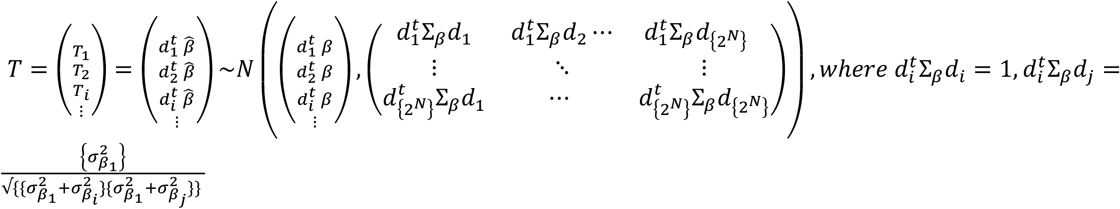 Under 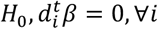 and we assume that the effect of all genotype combinations has the same distribution, i.e, under 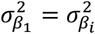. *So* under *H*_0_,

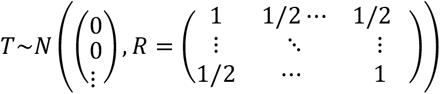 The p-value from the Max-T test can be computed using the following:

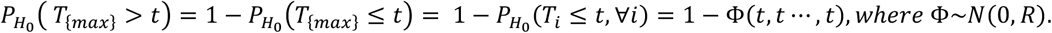 Max-T tests are conducted for both the epistasis SNP sets and the random N-way SNP collections (that are drawn n times). The p-values from the Max-T test provides an indication of how extreme the phenotypic trend of some combination is for the cluster tested. Step 3: Compute p-values for the epistasis set and the N-way SNP collections (n times generated) and check if: #p-values(Random SNPs groups) < p-value(Epistasis set). The Interactions Sets from the Interaction Score Matrix (*I*) in Supplementary Figure 3b are subjected to Max-T test (or anova test, see below) and the final interaction sets (that can be validated with Max-T or anova test) are stored in Interaction Score Matrix (*F*).

Often the phenotype distribution is not balanced amongst the different genotype combinations arising from the interaction of the N-SNPs. By construction, the genotype combination with all major alleles is expected to have a higher number of observations compared to the other genotype combinations. To ensure that the Max-T test is not influenced by the imbalance in the phenotype-genotype combination we also tested a Bootstrap version of the above Max-T test. We include the R code implementation of the Bootstrap version of the Max-T test in our GitHub repository.

#### Anova test

To make the statistical validation test more robust we also perform the anova test apart from the Max-T test and claim that an interaction is validated if it is performing well in at least one of the two tests. We perform the anova test for the Age 1 and Aging dataset using the following:

Age1 Dataset: For each N-way SNP set compute and evaluate the anova test using the following:

- Model1: *Y*~∑*SNP_i_* + *ϵ*, where *i* ∈ {1,… *N*}, *SNP_i_* are the SNPs in the N-way SNP interaction set.
- Model2: *Y*~∑*SNP_i_* + ∑*Genotype_j_* + *ϵ*, where *j* ∈ {1,… 2^*N*^}, *Genotype_j_* denote the different genotype combination arising from the interaction of N-SNPs.
- Anova(Model1, Model2) tests if Model2 is significantly better than Model1.
For Aging dataset, the test remains similar with the following modification:

- Model1: *Y*~∑*age*: *SNP_i_* + *ϵ*, where *i* ∈ {1,… *N*}, *SNP_i_* are the SNPs in the N-way SNP interaction set.
- Model2: *Y*~∑*age*: *SNP_i_* + ∑*age*: *Genotype_j_* + *ϵ*, where *j* ∈ {1,… 2^*N*^}, *Genotype_j_* denote the different genotype combination arising from the interaction of N-SNPs and *age*:*SNP_i_* captures interaction between *SNPi* and age or time on the phenotype
- Anova(Model1, Model2) tests if Model2 is significantly better than Model1.

The p-value obtained from the above test is denoted by lm_pval in this article. The above approach is analogous to the regression-based-test^56,57^ where logistic regression or linear regression are used to assess the impact of SNP interactions on diseases or quantitative traits. They typically compare the saturated model (Model 2) that includes interactions against the reduced model (Model 1) that omits interactions using likelihood ratio tests. Note that while the Max-T test is scalable to higher order interactions, the anova test may not be always scalable to high order interactions. Since we obtain interaction of maximum order 4 in the DGRP dataset, we applied both anova and Max-T test here.

### Computation efficiency

The computational accuracy of the method is better illustrated with a figure (see Supplementary Figure 10) where we show the time taken to apply MEIF on datasets with number of predictors varying in between 20 to 200. The computations are done in an Intel(R) Core(TM) i7-8550U CPU @ 1.80GHz 1.99 GHz processor laptop with 16 GB RAM. MEIF implementations takes around a minute for a data with 200 causal SNPs, as evident in Supplementary Figure 10. Note that the run time depends partly on the parameter settings of the cforest. For the simulations and for Supplementary Figure 10, we made sure that the number of trees (ntree) is same as the number of predictors in the model. From our simulations and from the real data application, we observed that for large dataset (>500 predictors), we did not gain much by increasing the ntree parameter. Hence, we keep the ntree fixed at 500 if the number of predictors (or SNPs) is over 500.

### DGRP Cardiac Dataset

We validated the applicability of our approach on both cross-sectional and longitudinal datasets by applying *epiMEIF* on 1) A cross-sectional dataset: Cardiac performance at 1 week (Heartperiod) and 2) a longitudinal dataset: Cardiac aging (Heartperiod at 1-week and 4-week).

The epiMEIF method has been developed in the framework of a project aiming to identify the genetic architecture of natural variations associated to cardiac aging in Drosophila. We analysed the cardiac performances in a natural population of young (1 week) and old (4 week) flies from the Drosophila Genetic Reference Panel (DGRP^21^), a population consisting of 205 inbred lines derived by 20 generations of full-sib inbreeding from inseminated wildtype caught female flies from the Raleigh, USA population. Whole-genome sequencing data, along with genotype calls, are available for all 205 lines. Contractility and rhythmicity were measured in females, using approximately 2000 flies from 168 DGRP lines at 1 week and 1800 flies from 165 lines at 4 weeks. More details on the phenotype and genotype dataset can be found in Saha et al.,^25^. Here, we have mainly focused on the application of *epiMEIF* on the natural variations of the Heart period from the above dataset. To alleviate the computation burden, we apply a single locus association test using mixed model (LMM) that first filters the SNP markers that have a significant effect on the phenotype, thereby reducing the size of the genome data for epistasis detection using epiMEIF. Details on data pre-processing steps involving the quality control checks on the DGRP GWAS dataset, followed by the single GWAS application is detailed below. An additional dataset of 20 DGRP lines was also analysed by us where the cardiac performance was studied following the same approach (with 12 flies observed per line) that were not used in the model building stage. We treat this dataset as our validation dataset and utilized it to test if the genetic interactions predicted from *epiMEIF* method in the original dataset have an impact on the phenotypic variations in this independent dataset.

#### Pre-processing steps and single GWAS

Genetic variants/genotype data linked to the cardiac phenotypes were identified using the GWAS web tool developed for the DGRP by Mackay^21^ (http://dgrp2.gnets.ncsu.edu/). It comprised four million high-quality SNPs amongst which 95% were homozygous. Typical quality assurance procedures involving maintaining the genotype call (GC) rate at 90% for each chromosome, removing SNPs with minor allele frequency (MAF)>0.0179 (must be present in 3 lines out of 167) were performed to get rid of the poor-quality genotype data. All phenotypes’ distributions are treated for outliers using standard outlier treatment i.e., remove the observations that lie outside 1.5 * IQR, where IQR, the ‘Inter Quartile Range’ is the difference between 75th and 25th quartiles. The removed data were validated by the biologists as technical outliers by looking back at the original experiment results. After the QC checks, single marker GWAS using LMM was performed on 2,456,752 SNPs from the DGRP data. The rapid decline in LD locally and lack of global population structure in Drosophila was favourable for GWAS application^21^. As mentioned in the main text, the single GWAS act as a filtering step and help narrow the search down to a reduced set of causal SNPs to accommodate computational requirements of *epiMEIF*. The nominal significance threshold of 10^−5^ was used to select the SNPs associated with the phenotype^21^. Note that if the number of causal SNPs (that needs to be explored for epistasis) is less than 10000, any filtering step can be avoided as the forest algorithm can handle as many as 10000 markers. Apart from the genome data, the DGRP dataset comprised other covariates like *Wolbachia pipientis* status that is a maternally transmitted endosymbiotic bacterium that infects ~53% of the DGRP lines and often impacts the phenotype of the flies^58,59^. These covariates can be easily accounted for in *epiMEIF*. The proposed method can simultaneously capture several vital components of genome-wide datasets: 1) The joint effect of the multiple loci via the function *f* in *equation*(2). 2) The interaction effects between the variants and in between the variant and “Time” (age in the DGRP dataset) (via the function *f* in *equation(2)*). 3) The effect of other covariates like Wolbachia pipientis status or population structure in the data using the covariate X in *equation(2)*. 4) the random effects due to the line/strain (with the help of covariate **Z** in *equation (2)*).

Results from GWAS: Subsequent to single-marker GWAS (via LMM) application, the SNPs significant at significance threshold 10^−5^ were tested for LD correction. Firtly, the SNPs were mapped to genes using the following rules: (i) if the variant is located in a gene (INTRON, EXON, 3’or 5’ UTR), the SNP is associated with this gene, (ii) if not, the variant is associated with a gene if its distance from the boundary of the gene (Transcription Start Site (TSS) and Transcription End Site (TES) is lesser than 1kb (iii) if no previous rules was applied, the variant is not mapped to gene. Then the SNPs that are highly correlated (above 90%) and that are the in the immediate neighbourhood of 5 SNPs, which are mapped to the same gene are clustered into the same LD blocks. Only one SNP (least causal/significant) is selected from each LD block for epiMEIF application. This led to 3484 SNPs for the cardiac HP dataset at 1 week and 1682 SNPs for the cardiac aging dataset.

### Simulation Study

We used genetic features obtained in the form of SNPs from the DGRP. For each simulation experiment, we considered a random subset of 1,000 SNPs with a minor allele frequency (MAF) >0.1. Using the MAF frequency of each SNP we artificially generated a dataset of 1000 SNPs using Bernoulli distribution and made sure that there is no population structure in the genotype dataset generated. We retained the same phenotype data (Heartperiod) as in the original cardiac dataset where there were 165 lines of flies and each line had approximately 10-12 flies. We wanted to show the effectiveness of our method for both longitudinal and cross-sectional data, so we designed two different simulation models: one for 1-week flies and another for both 1 week and 4-week flies.

To simulate the phenotype for week 1 flies, 6-8 SNPs were randomly drawn amongst which four chosen SNPs were considered as causal markers to simulate linear additive effects and further 1 to 3 pairs of SNPs were chosen to contribute epistatic effects. This is shown in Model 1. To simulate the aging phenotype, we use the model shown in Model 2 where the effect of the SNPs at age 4 weeks is higher than effect of the SNPs at age 1 week.

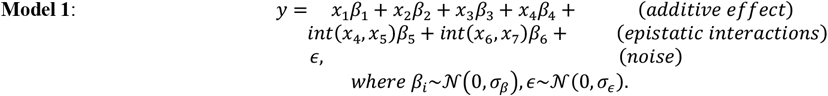

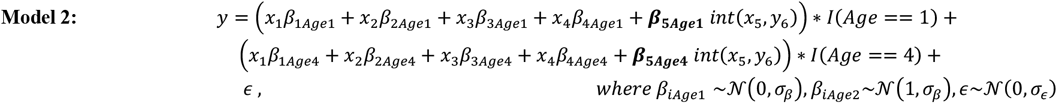

Interaction effects were simulated by taking the component-wise binary product as indicated by the ‘int’ operator above. This corresponds to interactions between both minor alleles. For the sake of simplicity, we did not consider other interactions (minor–major, major–minor and so on). The resulting vector is multiplied by the simulated effect size *β_i_*. The above model and its parameters are mainly adapted from Stephan et al.,^60^. However, unlike them, we do not consider any parameter separately for capturing the population structure because there is no underlying population structure in our artificial dataset. *σ_β_* and *σ_ϵ_* are assumed to be 0.375 and 0.125 respectively following Stephan et al.,^60^. To compare the performance of our method in different genetic settings, we vary the relative number of additive and interaction terms.

The goal of the simulation experiments is to check how accurately the *epiMEIF* can capture the above desired interactions in the different scenarios from the artificial dataset with simulated phenotype and artificially generated genotype data. Apart from the 6-8 SNPs shown above, we randomly choose 30 and 50 other SNPs (from the set of 1000 SNPs) in our simulation scenarios. To make sure that these additional SNPs used in the cforest do not have any marginal effect on the phenotype we ran a single marker GWAS model on the 1000 SNPs and extracted only those SNPs that were not associated with the phenotype. Amongst these SNPs, we selected 30 or 50 SNPs for our simulations. We conducted 100 simulations for each scenario and for each simulation we run the MEIF 10 times and obtain the interactions and their pooled interaction score from the 10 forests. We fit both the *epiMEIF* and the *Weighted epiMEIF* on the simulated cross-sectional data and only *epiMEIF* in longitudinal dataset. The *Weighted epiMEIF* was introduced to test how accurately they can capture the interactions around the SNPs that are marginally less significant in the model.

Apart from testing the overall power (see Table 1) of detecting the true interactions we also tested how often the true interactions are captured as the top-ranking interactions (based on the score) in our simulation scenarios. To evaluate this for each simulation, we rank the interactions obtained from the random forest application based on their score and compute the proportion of times the true interactions are captured across the different ranks. We call it the rank-wise power detection across the different simulation scenarios. Supplementary Figure 1 and Supplementary Figure 2 shows the rank-wise power in capturing the interaction of interest for Age 1 and Aging simulations, respectively. Supplementary Figure 1 illustrates that the *Weighted epiMEIF* can more efficiently capture the true interaction compared to the *epiMEIF*. Hence, we prefer *Weighted epiMEIF* for epistasis detection in cross-sectional data. Note that the improvement of *Weighted epiMEIF* over the *epiMEIF* is more prominent amongst the top ranks.

As already mentioned earlier, for aging data simulations, we have only applied *epiMEIF* and it performs very well in capturing the interactions with high marginal effect in the top ranks. Though less efficient in capturing interactions with low marginal effect (SNP_1: SNP_2 in Scenario 2 of Supplementary Figure 2), the overall power of capturing these interactions still reaches 80-90%.

### Forest parameter settings

In general, we aimed to keep parameter settings MEIF consistent and comparable across all scenarios. For the simulation scenarios, the random forest was built using 50 trees (ntree=50) and a random subsample of 1/3 (mtry=#No of predictors/3) of all available predictors were used to determine the best split. It was set at the default setting of random forest when categorical covariates are used in the model. ntree was chosen after model tuning.

Following other articles on simulations with epistatic data^12^, the number of simulations for each scenario was restricted to 100. The model efficacy did not improve much above this threshold. Note that the simulations scenarios are built using linear regression model with normality assumption. Splitting of tree nodes was stopped if they contained less than five samples. For the real DGRP data, we set the number of trees at 500 (chosen after model tuning).

## Acknowledgments

We thank Fabiana Rossi for her assistance on data analysis.

This work was supported by a research grant from Excellence Initiative of Aix-Marseille University - A*MIDEX (A*Midex International) to L.P. and a postdoctoral fellowship from the Turing Center for Living Systems (CENTURI) to S.S.

## Supplementary Tables

**Supplementary Table 1:** Proportion of 2-way, 3-way and 4-way interactions that can be validated from DGRP HP data (1 week) with the validation cohort.

| *epiMEIF* interactions | #2-way | #3-way | #4-way | Total |
| --- | --- | --- | --- | --- |
| Total | 45 | 33 | 2 | 80 |
| Not eligible for testing with the validation data due to lack of data | 8 | 2 | 0 | 11 |
| Showing effect with validation data | 5 | 13 | 2 | 20 |
| % Interactions validated | 14%(5/37) | 42%(13/31) | 100% (2/2) | 29%(20/69) |

**Supplementary Table 2:**

**Summary of the list of genetic variants and their biological significance that are involved in the statistical interaction network obtained by applying epiMEIF on DGRP Heart period at 1-week data.**

**Supplementary Table 3:**

**Summary of the detailed list of interactions along with their scores and statistical significance from the epiMEIF network on DGRP Heart period at 1-week data.**

**Supplementary Table 4:**

**Summary of the list of genetic variants and their biological significance that are involved in the statistical interaction network obtained by applying epiMEIF on DGRP Heart period during aging data.**

**Supplementary Table 5:**

**Summary of the detailed list of interactions along with their scores and statistical significance from the epiMEIF network on DGRP Heart period during aging data.**

**Supplementary Table 6:**
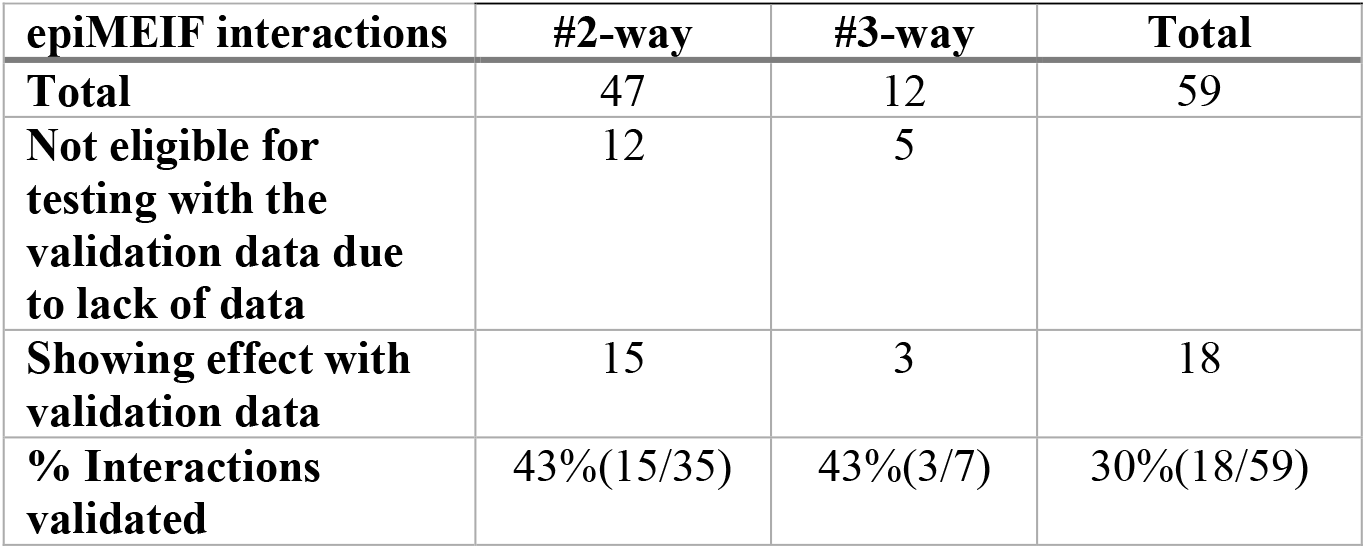
Proportions of 2-way and 3-way interactions that can be validated from DGRP Heart period data during aging with the validation cohort.

## Supplementary Figures Legends

**Supplementary Figure 1:**
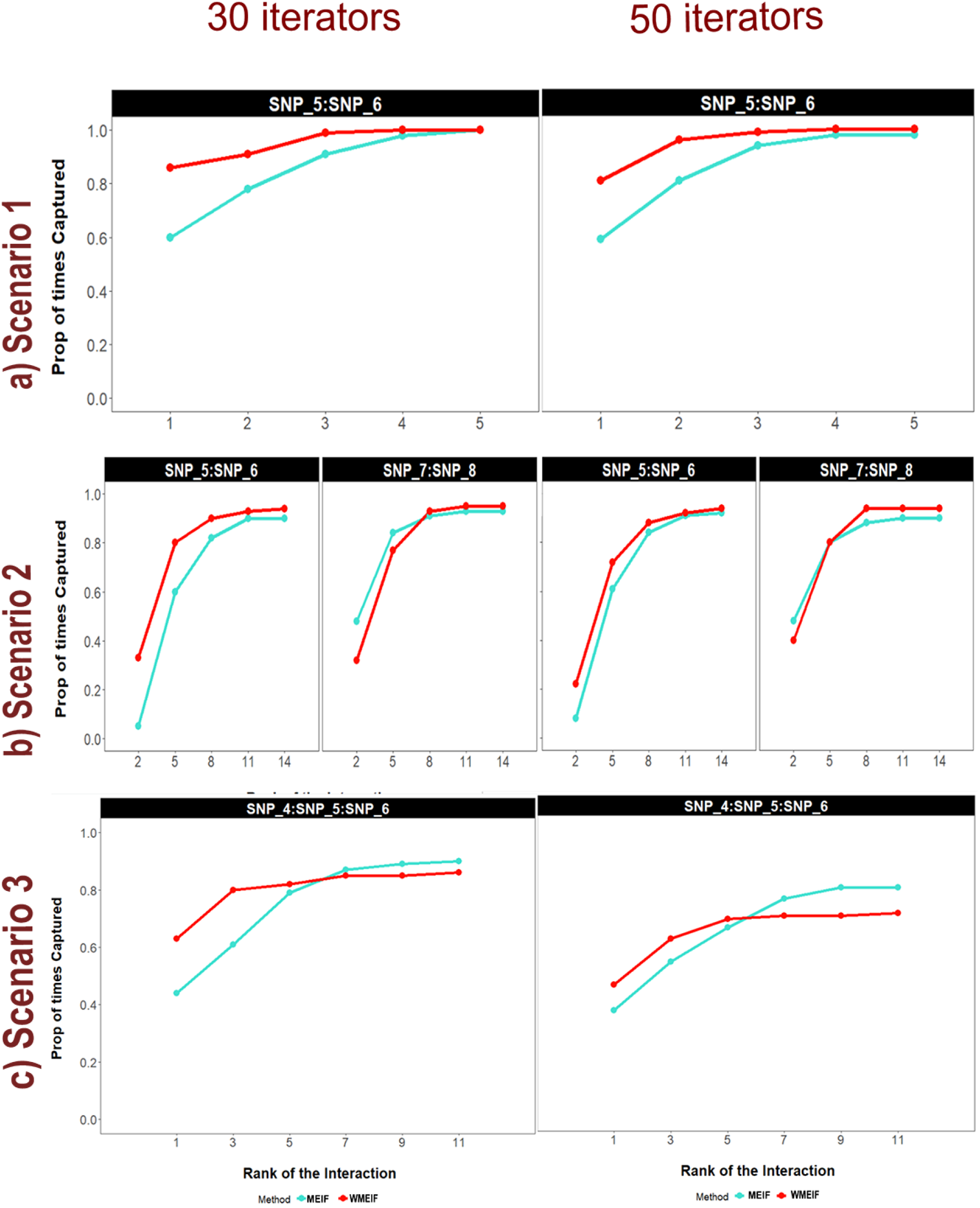
Overall power in detecting the true interactions in the simulation scenarios S1, S2 and S3 across the different ranks. For each scenarios the frequency at which true model interactions are captured in the top n rank interactions is displayed (with 30 (left) or 50 (right) iterators). *n* is so chosen such that it can capture the minimal and maximal optimum power of interaction in the above simulation scenarios.

**Supplementary Figure 2:**
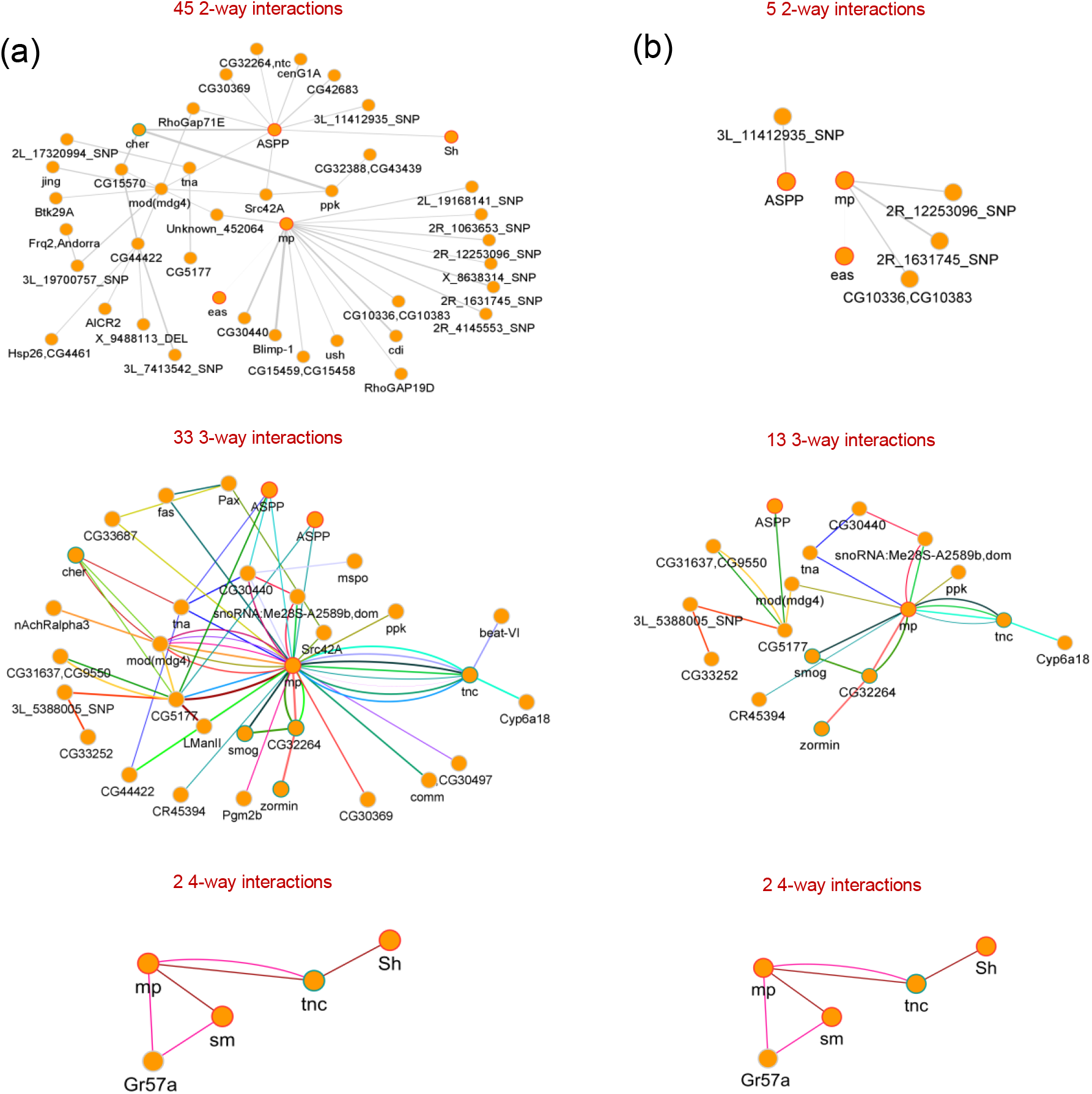
(a) Different n-way interactions (n=2, 3 and 4) obtained when weighted MEIF is fitted on DGRP Heart period 1 week training dataset. Higher order interactions (more than binary) are highlighted with different coloured edges. (b) Different n-way interactions from part (a) that can be validated in the validation dataset using anova/Max-T test.

**Supplementary Figure 3:**
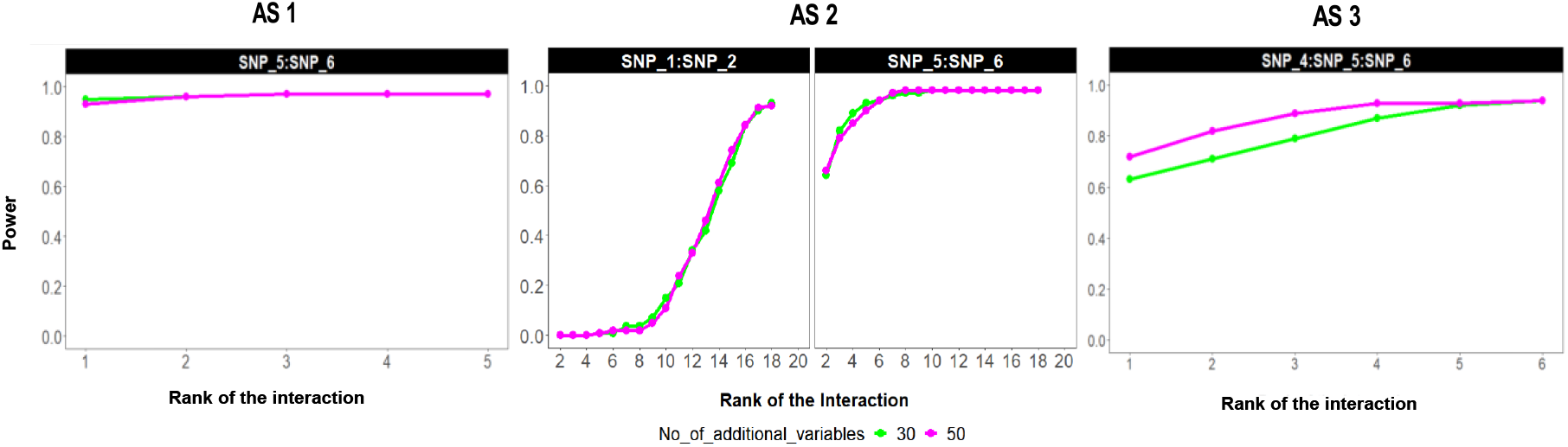
The figure shows the power of detecting the true interactions in simulation scenarios AS1, AS2 and AS3 across the different ranks. For each scenario, the plot shows how often the true model interactions are captured in the top n ranks, where n varies between 1 and 5 for AS1, n varies between 2 and 22 for AS2 and n varies between 1 and 10 for AS3. n is chosen to capture the minimal and maximal optimum power of interaction in the above simulation scenarios. The two different coloured curves indicate the two different simulation scenarios where the number of additional non-causal SNPs used in the model is 30 (green curve) and 50 (magenta curve).

**Supplementary Figure 4:**
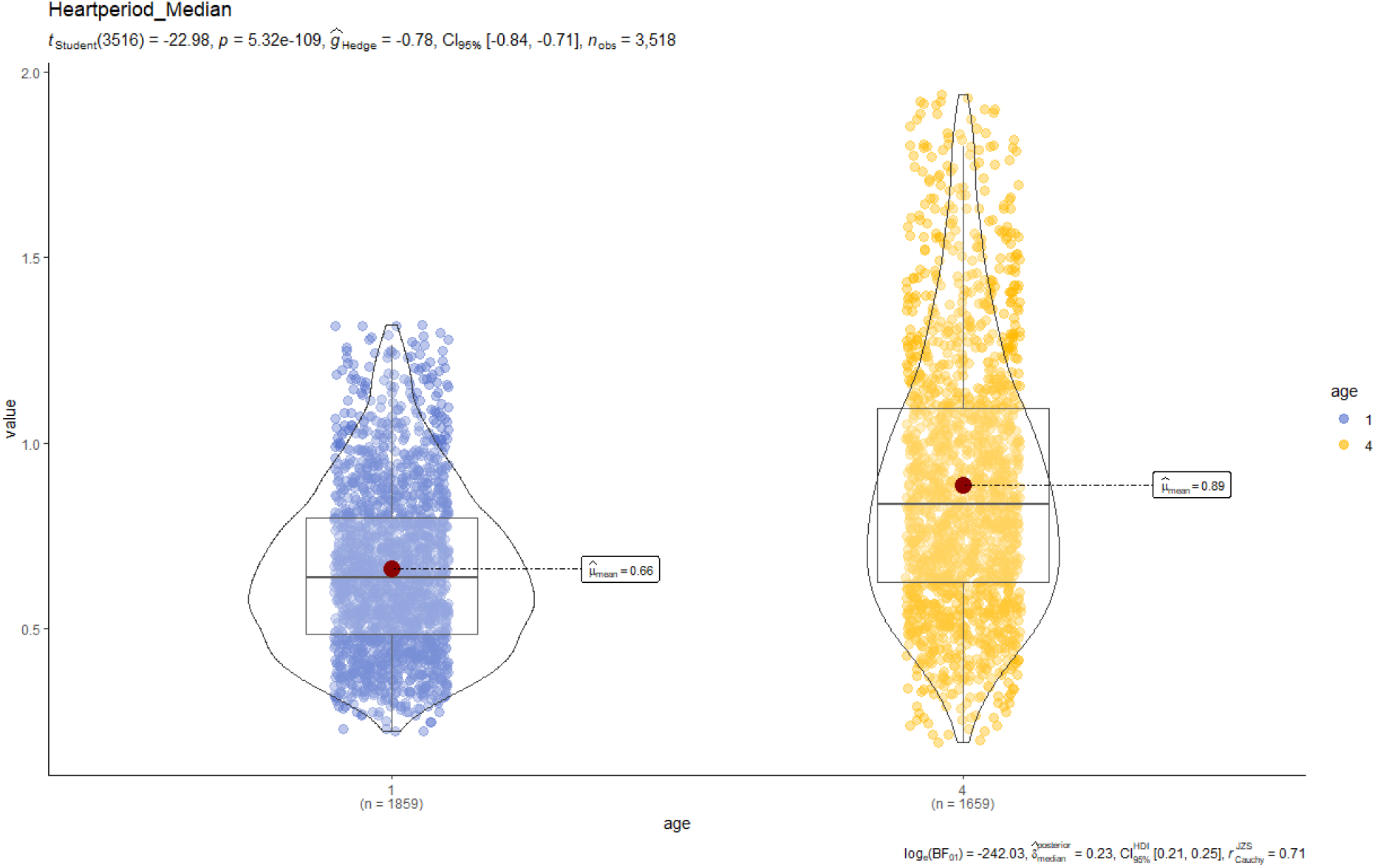
Significant Change of Heart Period in the DGRP from 1 week to 4 weeks

**Supplementary Figure 5:**
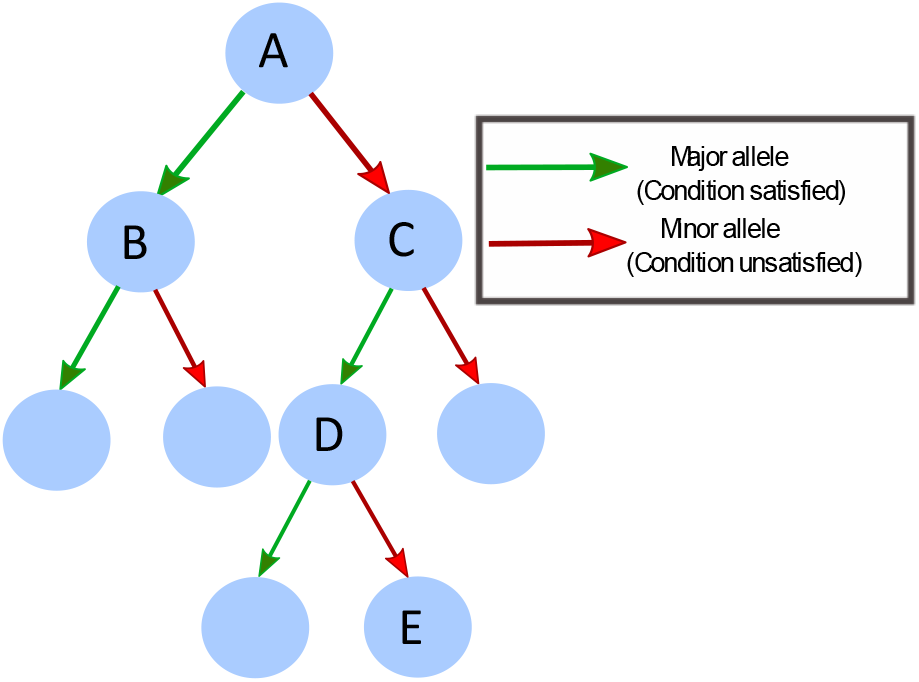
Example tree generated by the cforest algorithm. SNP pairs A and B, A and C, A and D, A, and E, C and D, C and E, and D and E represent descendant pairs, which may indicate epistatic genetic effects, whereas SNP pairs B and C, B and D, and B and E represent non-descendant pairs, which may indicate independent additive genetic effects.

**Supplementary Figure 6:**
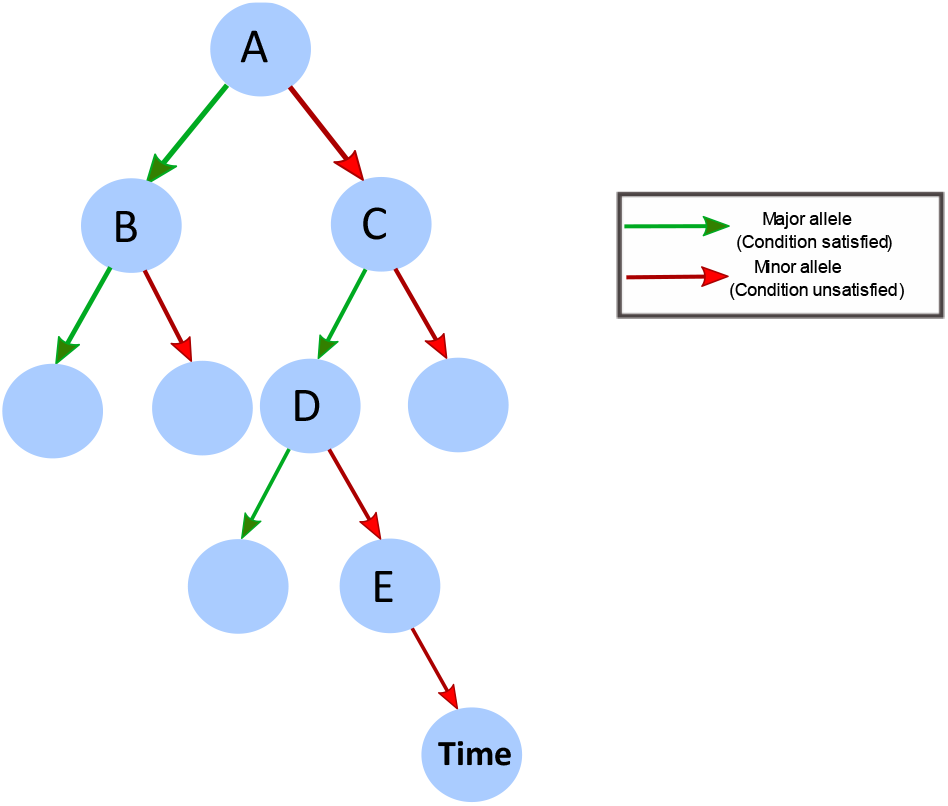
Example tree generated by the Random Forests algorithm when applied on a longitudinal dataset.

**Supplementary Figure 7:**
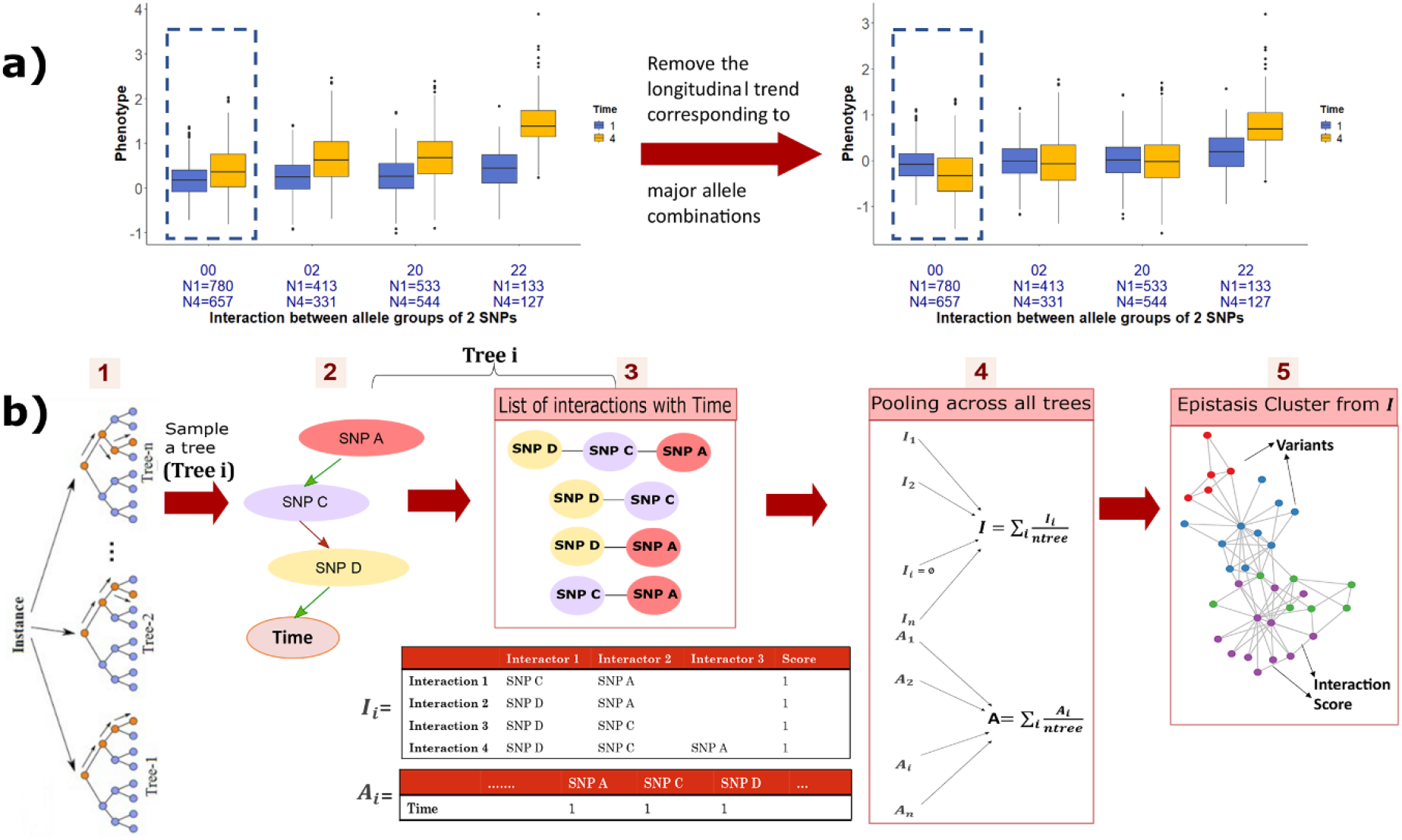
a) Boxplot distribution of cardiac phenotype from the Drosophila Cardiac Genetics Dataset across different genotype combination arising from the interaction of 2 SNPs. The longitudinal phenotypic trend corresponding to the major allele combinations (00) removed from the overall data prior to the exploring the epistatic interactions in the dataset. b) gives a step-by-step illustration of how epistasis clusters are formed from the trees in the random forest.

**Supplementary Figure 8:**
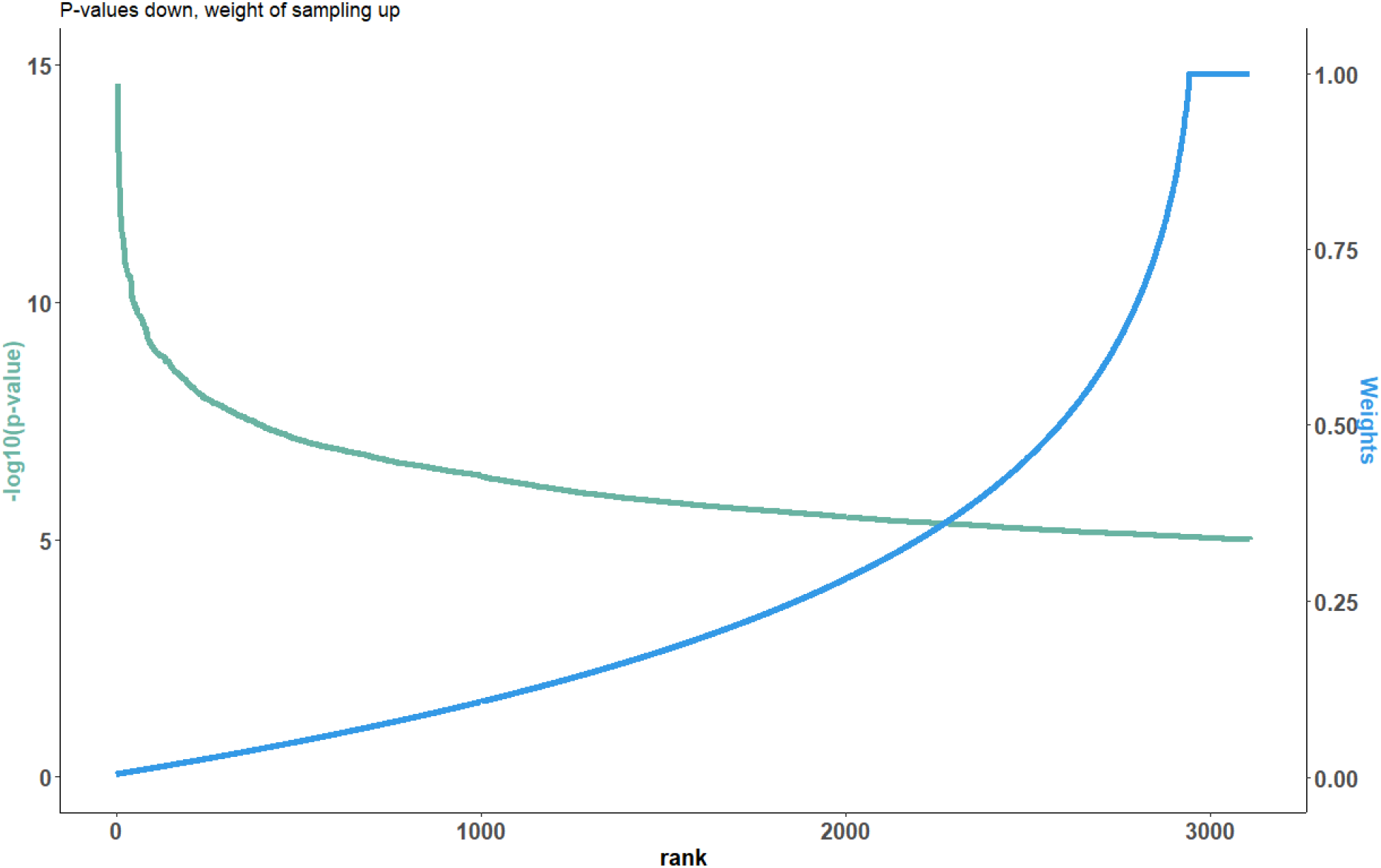
The figure plots the −log10(pvalues) of the SNPs from the single marker GWAS on the Heart period variation of the DGRP dataset (at 1 week). The SNPs are arranged based on their ranks (most significant getting the lowest rank). The secondary axis here shows the weights assigned to each SNP in the weighted random forest. The weights are so designed, such that they are inversely proportional to the pvalues.

**Supplementary Figure 9:**
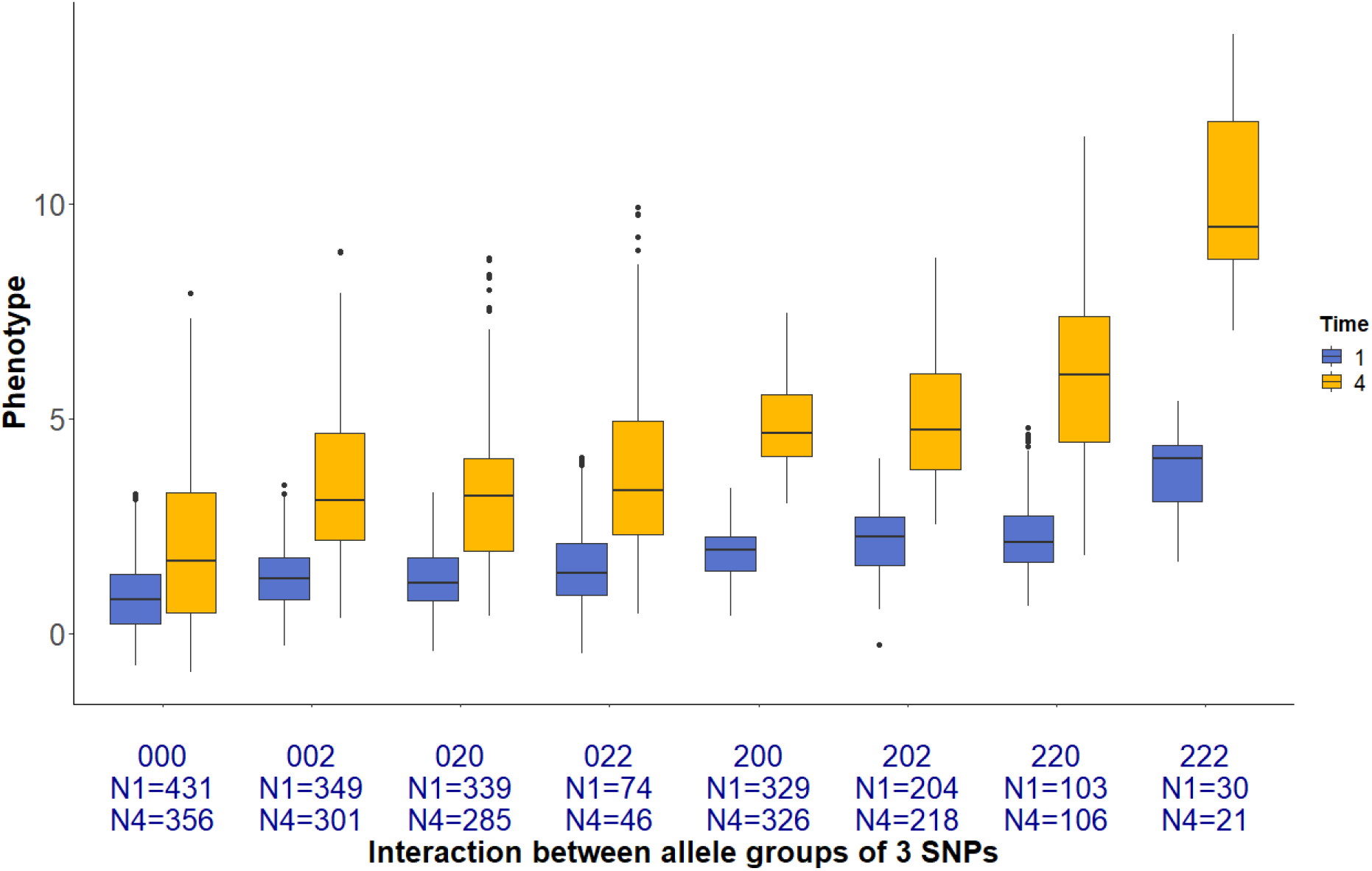
Boxplot distribution of the cardiac phenotype “Heart period” of the DGRP flies across different genotype combination arising from the interaction of 3 SNPs. Within each allele combination the boxplot distribution of the phenotype of flies at 1 week and at 4 weeks are highlighted by blue and yellow. The observation sample size within each population and in each genotype combination is shown in the x axis

**Supplementary Figure 10:**
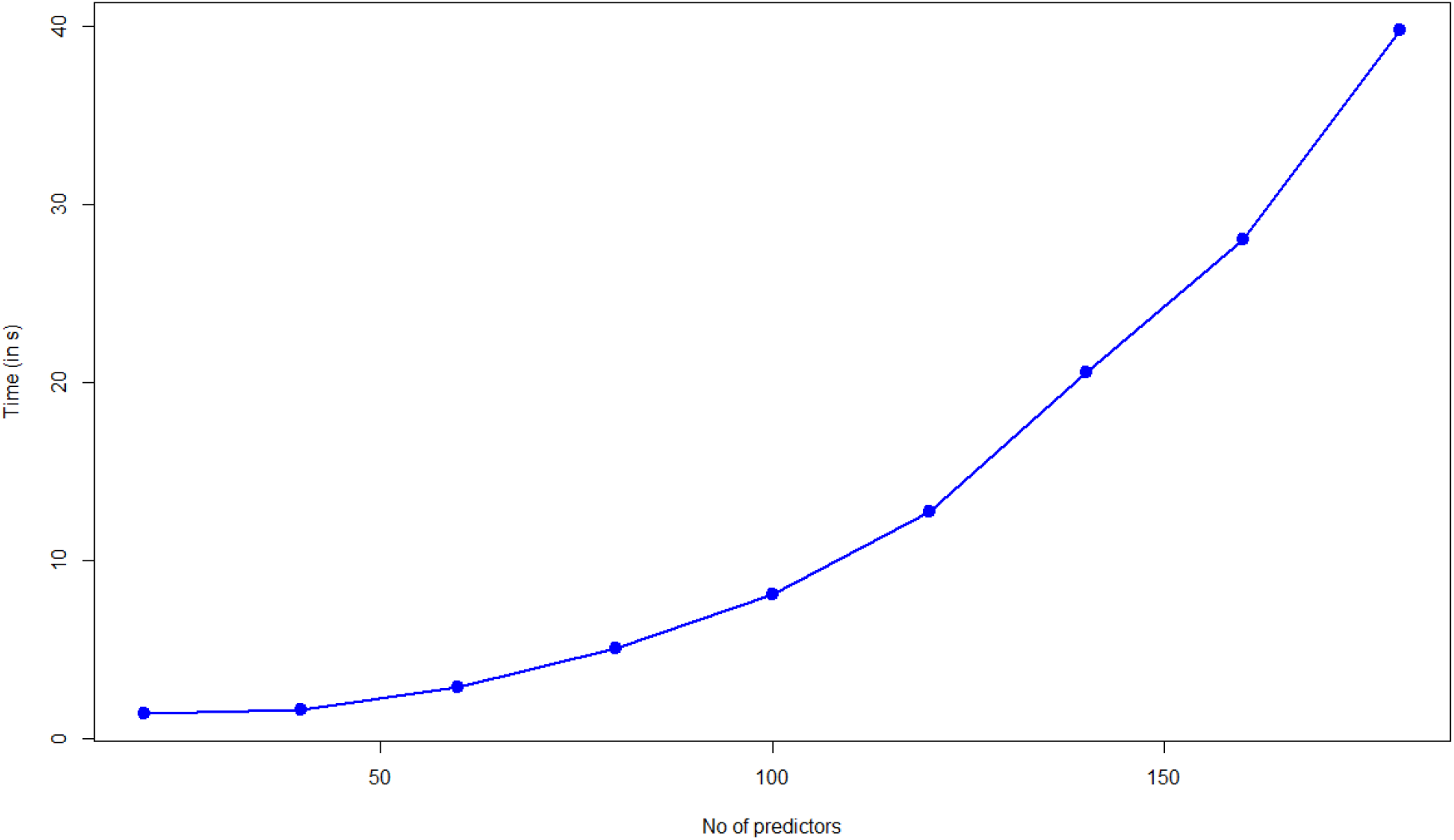
Computational efficiency of epiMEIF: time taken to apply MEIF on datasets with number of predictors varying in between 20 to 200.

